# The PDIM paradox of *Mycobacterium tuberculosis*: new solutions to a persistent problem

**DOI:** 10.1101/2023.10.16.562559

**Authors:** Claire V. Mulholland, Thomas J. Wiggins, Jinhua Cui, Catherine Vilchèze, Saranathan Rajagopalan, Michael W. Shultis, Esmeralda Z. Reyes-Fernández, William R. Jacobs, Michael Berney

**Affiliations:** Department of Microbiology and Immunology, Albert Einstein College of Medicine, Bronx, New York, USA

## Abstract

Phthiocerol dimycocerosate (PDIM) is an essential virulence lipid of *Mycobacterium tuberculosis*. *In vitro* culturing rapidly selects for spontaneous mutations that cause PDIM loss leading to virulence attenuation and increased cell wall permeability. We discovered that PDIM loss is due to a metabolic deficiency of methylmalonyl-CoA that impedes the growth of PDIM-producing bacilli. This can be remedied by supplementation with odd-chain fatty acids, cholesterol, or vitamin B_12_. We developed a much-needed facile and scalable routine assay for PDIM production and show that propionate supplementation enhances the growth of PDIM-producing bacilli and selects against PDIM-negative mutants, analogous to *in vivo* conditions. Our results solve a major issue in tuberculosis research and exemplify how discrepancies between the host and *in vitro* nutrient environments can attenuate bacterial pathogenicity.

## Main Text

The cell wall of *Mycobacterium tuberculosis* (*Mtb*) is exceptionally complex and is essential to its success as a pathogen. Phthiocerol dimycocerosates (PDIMs) are long-chain non-polar lipids found in the outermost layer of the cell wall of *Mtb* and other pathogenic slow-growing mycobacteria^1^. PDIMs play a crucial role in *Mtb* pathogenesis (reviewed in^2^), however, *Mtb* is prone to losing the ability to produce PDIM *in vitro* due to spontaneous mutation of PDIM biosynthesis genes^3,4^. Loss of PDIM biosynthesis confers a growth advantage in current mycobacterial culture media^3,5^, resulting in PDIM-deficient mutants dominating cultures with successive passage^3^. As PDIM deficiency decreases virulence^5–11^ and increases cell wall permeability^12,13^, spontaneous PDIM loss adversely affects experimental reliability, reproducibility, and the interpretation of results. PDIM deficiency has also been shown to reduce the vaccine efficacy of *Mycobacterium bovis* BCG Pasteur^14^. “The PDIM problem” thus presents a major challenge in tuberculosis research and has hindered progress in the field for decades. We sought to understand the underlying cause of PDIM loss and develop routine methods to enable reproducible PDIM bias-free investigations in all branches of *Mtb* research.

### A tractable and scalable PDIM screen

The genetically unstable nature of the PDIM biosynthetic pathway makes routine PDIM screening essential for all branches of tuberculosis research. However, current PDIM screening approaches such as whole genome sequencing (WGS), mass spectrometry, and thin layer chromatography (TLC), are expensive, cumbersome, and require specialized equipment and expertise, further compounding the PDIM problem.

We hypothesized that the differential permeability of PDIM-positive [PDIM(+)] and PDIM-negative [PDIM(-)] *Mtb*^12,13^ could be exploited to develop a simpler functional PDIM assay. To test this, we first assembled a PDIM reference strain set comprised of six BSL2-approved attenuated *Mtb* H37Rv strains with varying PDIM content (Fig. 1a, Supplementary Table 1). These strains demonstrate the heterogeneity of PDIM production commonly found in laboratory *Mtb* strains. Vancomycin – a large antimicrobial glycopeptide not normally used for *Mtb* treatment due to poor penetration, has previously been reported to be more effective against PDIM deletion mutants of *Mtb* and *M. bovis* BCG than the corresponding PDIM(+) wildtype strains^15^. Accordingly, we found that PDIM levels measured by TLC significantly correlated with vancomycin MIC_90_ and MIC_50_ in our reference strain set after 10–14 days of incubation (Supplementary Fig. 1). PDIM(+) *Mtb* mc^2^7902 was also more resistant to other high molecular weight compounds than PDIM(-) mc^2^8398, though vancomycin gave the best differentiation (Extended Data Fig. 1a,g). Furthermore, much greater Ethidium Bromide uptake^16^ was observed in mc^2^8398 than mc^2^7902 (Extended Data Fig. 1h), consistent with enhanced permeability of PDIM(-) strains.

**Fig. 1.**
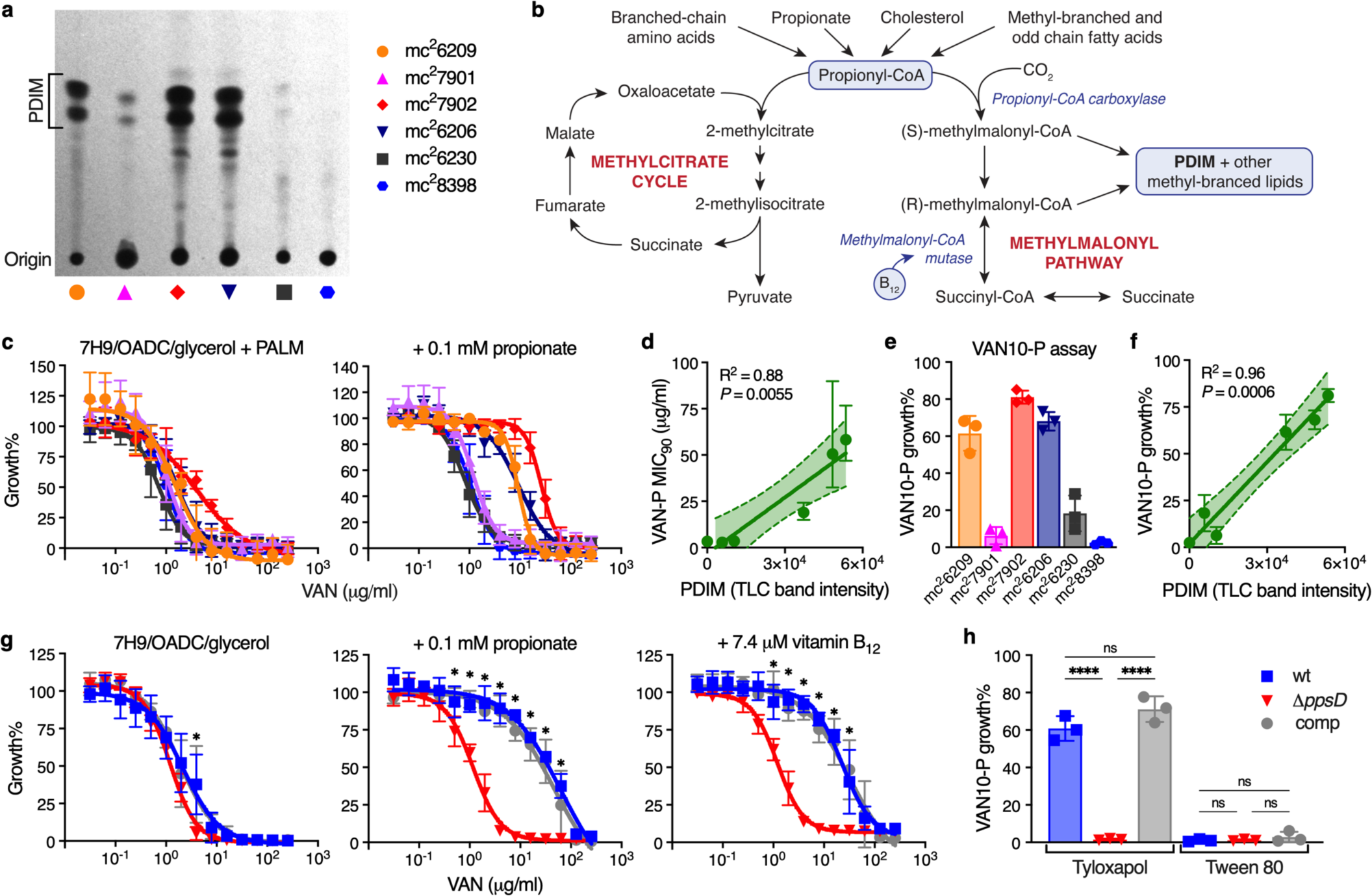
| Vancomycin resistance is enhanced by propionate or vitamin B_12_ supplementation and is predictive of PDIM production in *Mtb*. **a**, TLC lipid analysis of the PDIM reference strain set (see Supplementary Table 1). **b**, Metabolic pathways of methylmalonyl-CoA production and propionyl-CoA catabolism. **c**, Vancomycin resistance of the *Mtb* PDIM reference strain set in 7H9/OADC/glycerol/tyloxapol + PALM (pantothenate, arginine, leucine, and methionine) media, and additionally supplemented with 0.1 mM propionate (‘VAN-P’ MIC), measured after 7 days incubation. **d**, Correlation between VAN-P MIC_90_ ± 95% CI from the curve fit in (**c**) (+ 0.1 mM propionate) and PDIM band intensity from (**a**). The band represents the 95% CI. **e**, ‘VAN10-P’ assay comparing growth in 10 μg/ml vancomycin with 0.1 mM propionate to drug-free controls (VAN10 OD / VAN0 OD ξ 100 = VAN10 growth%). **f**, Correlation between VAN10-P growth% from (**e**) and PDIM from (**a**). **g**, Vancomycin resistance of isogenic PDIM(+) and PDIM(-) *Mtb* H37Rv strains in standard 7H9/OADC/glycerol/tyloxapol media and supplemented with 0.1 mM propionate or 7.4 μM vitamin B_12_ (10 μg/ml). \**P* < 0.001 for both wt and comp versus 11.*ppsD*; two-way ANOVA with Tukey’s multiple comparison test. **h**, VAN10-P assay of H37Rv strains with tyloxapol or Tween 80. \*\*\*\**P* < 0.0001; one-way ANOVA with Tukey’s multiple comparison test. MIC data show mean ± SD for *n* = 4 biological replicates from two independent experiments. VAN10-P data show mean ± SD for *n* = 3 three independent experiments, each performed in triplicate.

Whilst an intact biosynthetic pathway is essential for PDIM production, PDIM size and abundance are dependent on the availability of the three-carbon precursor methylmalonyl-CoA (MMCoA)^17^. MMCoA is generated from propionyl-CoA by propionyl-CoA carboxylase, or, from succinyl-CoA by vitamin B_12_-dependent MMCoA mutase (Fig. 1b). In the host, *Mtb* has access to propionyl-CoA-generating carbon sources such as cholesterol^18,19^ and possibly also scavenges vitamin B ^20–22^. Standard Middlebrook 7H9/OADC/glycerol media, however, lacks both a propionyl-CoA-generating carbon source and vitamin B_12_. Propionate supplementation or growth on cholesterol as a sole carbon source have been shown to increase PDIM biosynthesis^17,23^. Accordingly, we found that the addition of 0.1 or 1.0 mM propionate preferentially increased vancomycin resistance of PDIM(+) strains in 7H9/OADC/glycerol/tyloxapol + PALM media (pantothenate, arginine, leucine, and methionine; for BSL2 auxotrophic strains), enhancing the differentiation between PDIM(+) and PDIM(-) *Mtb* while improving assay robustness and reducing time to result (Fig. 1c, and Extended Data Fig. 2). To further simplify our approach and enable scalability, we established a single concentration assay we term the ‘VAN10-P assay’ (Fig. 1e, Extended Data Fig. 2c and Supplementary Fig. 2). The VAN10-P assay compares growth in 10 μg/ml vancomycin with 0.1 mM propionate to no-drug controls and highly correlates with PDIM production (Fig. 1f). We additionally validated our approach using isogenic PDIM(-) (Δ*ppsD*) and complemented (Δ*ppsD*::comp) strains constructed from a PDIM(+) clone (H37Rv-SC, wildtype) (Supplementary Table 2). PDIM(+) and PDIM(-) H37Rv showed a greater than 30-fold difference in vancomycin MIC_90_ with 0.1 mM propionate (‘VAN-P’ MIC) (Fig. 1g) and this was also reflected in the VAN10-P assay (Fig. 1h). Highly similar results were also obtained for *Mtb* CDC1551 and its isogenic PDIM(-) (11.*mas*) mutant (Supplementary Fig. 3).

To confirm that increased vancomycin resistance with propionate was due to enhanced PDIM production rather than other effects such as accumulation of propionyl-CoA or methylcitrate cycle intermediates^24,25^, we supplemented with vitamin B_12_ to provide an alternate route for MMCoA production via the vitamin B_12_-dependent methylmalonyl pathway^22^ (Fig. 1b). Vitamin B_12_ selectively increased the vancomycin resistance of PDIM(+) *Mtb* mirroring the effect of propionate (Fig. 1g and Extended Data Fig. 2b,c), consistent with enhanced resistance due to increased PDIM production. Leucine, a potential source of propionyl-CoA^26^, did not have a marked effect on vancomycin resistance of H37Rv at the concentration provided in PALM- supplemented media (0.38 mM) (Supplementary Fig. 4a).

As Tween 80 is another detergent commonly used in *Mtb* culture media, we tested whether tyloxapol could be replaced with Tween 80 in our assay. Tween 80 is known to remove several layers of the mycobacterial cell wall including PDIM^27^. Consistent with this, Tween 80 abolished PDIM-related differences in vancomycin resistance and further increased the vancomycin sensitivity of PDIM(-) *Mtb* (Fig. 1h and Extended Data Fig. 3).

### Breadth and depth of PDIM bias in tuberculosis research

Next, we determined the predictive power of our approach in a range of virulent *Mtb* strains including Erdman, HN878, KZN 4207, and two different CDC1551 and H37Rv stocks (Supplementary Table 2). VAN-P screening reliably predicted PDIM levels as determined by TLC for all these strains (Fig. 2a and Extended Data Fig. 4a–d). Furthermore, VAN-P assays outperformed WGS at diagnosing PDIM deficiencies in heterogeneous populations. TLC and VAN10-P assays showed low PDIM levels in H37Rv-A and CDC1551-A (Fig. 2a), however, standard WGS variant calling failed to identify any PDIM mutations in these stocks, whilst an unfixed mutation was identified in Erdman (Extended Data Table 1). Low-frequency variant analysis subsequently identified putative PDIM mutations at ∼10–13% frequency in each of these stocks (Extended Data Table 2), indicating they comprise a mixture of different PDIM(-) mutants. Thus, WGS is a poor predictor of PDIM levels in mixed populations as these can comprise an array of different low-frequency PDIM mutations, which can be difficult to detect by WGS.

**Fig. 2.**
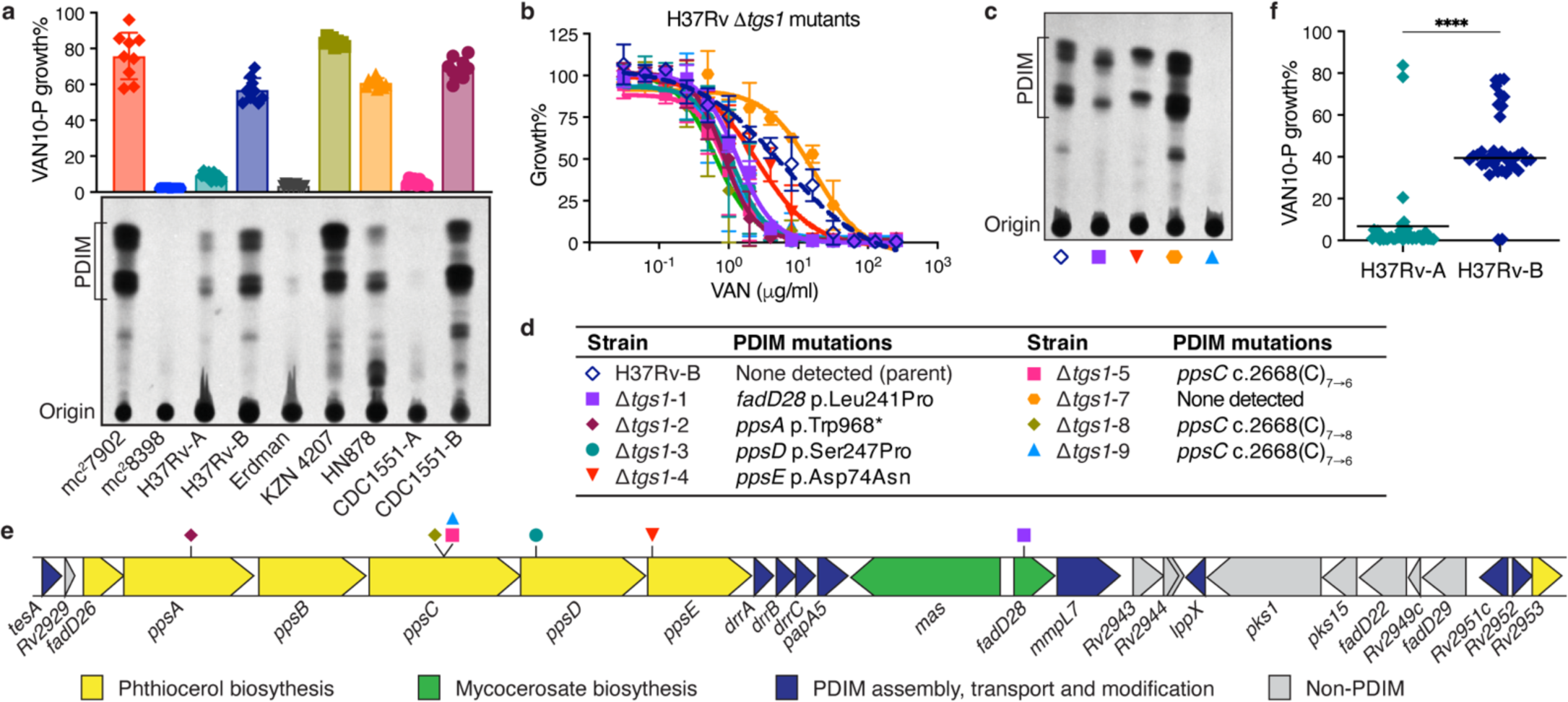
| VAN-P assays accurately predict PDIM status during genetic manipulations and across different *Mtb* strains and lineages. **a**, TLC lipid analysis and VAN10-P assays of different laboratory stocks of virulent *Mtb* strains alongside *Mtb* mc^2^7902 and mc^2^8398. Mean ± SD for *n* = 9 pairwise comparisons between triplicate wells. **b**, VAN-P MIC assays of eight 11.*tgs1* mutants and the parent H37Rv-B. Mean ± SD for *n* = 3–4 biological replicates from two independent experiments. **c**, TLC lipid analysis of four 11.*tgs1* mutants and H37Rv-B. Lipid extracts in (**a**) and (**c**) were run on the same TLC plate. **d**, Mutations in PDIM biosynthetic genes of 11.*tgs1* mutants (see also Extended Data Fig. 5). **e**, Schematic showing the PDIM gene cluster and location of secondary PDIM mutations in 11.*tgs1* mutants. **f**, VAN10-P screening of single colonies isolated from H37Rv-A (*n* = 38) and H37Rv-B (*n* = 37). Each colony was assayed in triplicate and data points represent mean VAN10-P growth%. Lines indicate the median. \*\*\*\**P* < 0.0001; unpaired two-tailed Mann-Whitney test.

To investigate how genetic engineering of *Mtb* strains is affected by PDIM loss we generated knockout mutants of the non-PDIM-related gene *tgs1* from a mouse-passaged H37Rv stock (H37Rv-B) using specialized transduction^28^. Surprisingly, despite this being a mouse-passaged stock, only one of eight 11.*tgs1* mutants obtained (11.*tgs1*-7) was found to be fully PDIM(+) by VAN-P MICs (Fig. 2b). This was further validated by TLC and sequence analysis (Fig. 2c–d). Historically animal passaging was the only procedure known to select for PDIM(+) *Mtb*. However, our specialized transduction results and VAN-P assays suggested that despite animal passage this may still be a mixed population (Fig. 2a–d). Indeed, VAN10-P single colony screening confirmed that while animal passage enriched for PDIM(+) clones, PDIM-deficient strains were not completely removed (Fig. 2f, Extended Data Table 1). Consequently, using VAN10-P single colony screening, we were able to isolate single PDIM(+) clones from H37Rv-B as well as other virulent strains and from avirulent mc^2^6230 (Extended Data Fig. 4 and Extended Data Table 1).

Strikingly, six different mutations in five different PDIM genes were identified across the seven PDIM(-) 11.*tgs1* mutants (Fig. 2d,e), emphasizing the genetic heterogeneity in the PDIM gene cluster in mixed populations. We also found two unique frameshift mutations in a 7-cytosine homopolymeric tract in *ppsC* (Extended Data Fig. 5). This region appears to be a ‘hotspot’ for mutation as we also found *ppsC* homopolymeric tract mutations in mc^2^6230 (Extended Data Fig. 5e) and in the literature^29,30^. Homopolymeric tracts are prone to mutations caused by slipped-strand mispairing^31^ and as *Mtb* lacks a DNA mismatch repair system^32^ this may lead to hypervariability in these regions, further augmenting the propensity for PDIM loss *in vitro*.

Collectively these data validate VAN-P assays as a reliable and effective method to assess PDIM levels and heterogeneity in *Mtb* populations and aid in the isolation of PDIM(+) clones. However, the data presented also strongly emphasized the need to resolve the underlying issue of PDIM loss.

### MMCoA deficiency impairs the growth of PDIM(+) *Mtb*

As PDIM production is tightly coupled to *Mtb* metabolism^17^, we reasoned that there may be a metabolic solution to the PDIM problem. Propionyl-CoA, an upstream precursor of PDIM, can be inhibitory to bacterial growth^33^. The major pathways for propionyl-CoA detoxification are the methylcitrate cycle^34^, the methylmalonyl pathway^22^, and the incorporation into PDIM and other virulence-associated lipids^35,36^ (Fig. 1b). We hypothesized that PDIM-deficient strains would be more sensitive to propionate toxicity without this sink for propionyl-CoA metabolism and that this could be exploited to create a PDIM selective medium. Consistent with this hypothesis, the PDIM(-) strains in our reference strain set were more sensitive to propionate than the PDIM(+) (Fig. 3a). Surprisingly, we also observed that PDIM(+) strains reached higher density at lower propionate concentrations (Fig. 3a), suggesting sub-toxic propionate may provide a growth advantage to PDIM(+) *Mtb*.

**Fig. 3.**
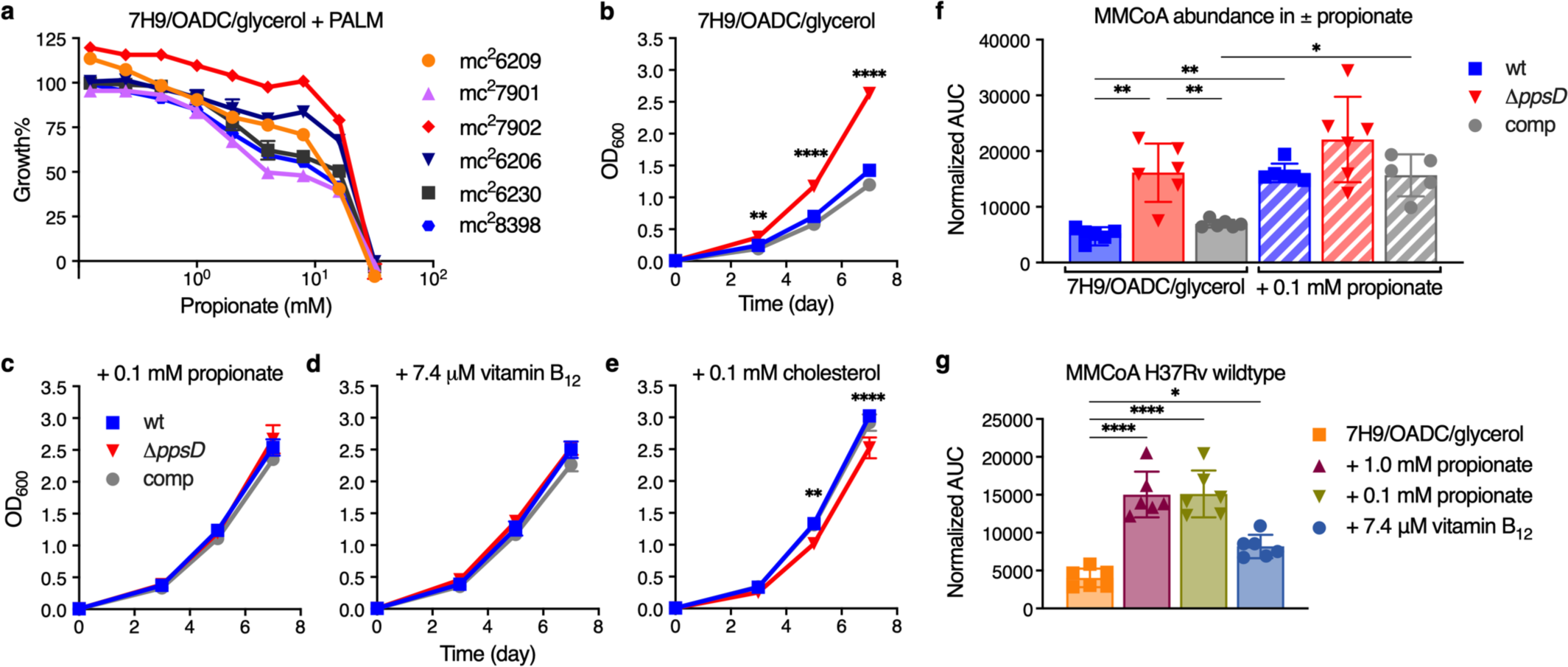
| Propionate and vitamin B_12_ supplementation restore the growth of PDIM(+) *Mtb*. **a**, Relative growth of the PDIM reference strain set in 7H9/OADC/glycerol/tyloxapol + PALM media with increasing concentrations of propionate compared to no propionate controls. Mean ± SD for *n* = 3 biological replicates. **b**–**e**, Growth curves of PDIM(+) and PDIM(-) *Mtb* H37Rv in (**b**) standard 7H9/OADC/glycerol/tyloxapol and (**c**) supplemented with 0.1 mM propionate, (**d**) 7.4 μM vitamin B_12_ (10 μg/ml), or (**e**) 0.1 mM cholesterol. Mean ± SD for *n* = 3 biological replicates. \*\**P* < 0.01, \*\*\**P* < 0.001, \*\*\*\**P* < 0.0001 for both wt and comp versus 11.*ppsD*; two-way ANOVA with Šidák’s multiple comparison test. Data in (**a**–**e**) are representative of at least two independent experiments. For some data points the SD is smaller than the data symbols. **f**, Abundance of methylmalonyl-CoA (MMCoA) in PDIM(+) and PDIM(-) H37Rv grown in standard 7H9/OADC/glycerol/tyloxapol media ± 0.1 mM propionate, and **g**, PDIM(+) H37Rv wildtype in standard media and supplemented with either propionate or vitamin B_12_. Abundances are shown as normalized area under the curve (AUC). Mean ± SD for *n* = 6 biological replicates from two independent experiments. \**P* < 0.05, \*\**P* < 0.01, \*\*\*\**P* < 0.0001; one-way ANOVA with Tukey’s multiple comparison test. Significant differences between ± propionate for each strain and between strains for each condition are indicated in (**f**), and compared to unsupplemented media in (**g**).

Next, we compared the growth of isogenic PDIM(+) and PDIM(-) *Mtb* with propionate and other supplements. Notably, we found that the addition of 0.1 or 1.0 mM propionate to standard 7H9/OADC/glycerol/tyloxapol media increased the growth rate of PDIM(+) strains to that of PDIM(-) (Fig. 3c and Extended Data Fig. 6b,c). Vitamin B_12_ also restored PDIM(+) growth analogous to propionate (Fig. 3d). Again, we did not observe comparable effects when tyloxapol was replaced with Tween 80 (Extended Data Fig. 6k–m), implying more extensive disruption of PDIM(+) growth in this detergent. The odd-chain fatty acid valerate also restored PDIM(+) growth, but not the even-chain fatty acids acetate or butyrate, or the three-carbon metabolite pyruvate (Extended Data Fig. 6g–j), demonstrating this effect is specific to propionyl-CoA generating carbon sources. Similar to vancomycin resistance assays, we again did not observe a comparable effect with leucine supplementation (Supplementary Fig. 4), suggesting that at this concentration leucine is preferably routed into anabolic pathways rather than catabolised as a source of propionyl-CoA. Supplementing with cholesterol not only restored PDIM(+) growth but also significantly reduced the growth of PDIM(-) *Mtb* (Fig. 3e). The growth reduction of PDIM(-) *Mtb* is likely related to the reduced ability to maintain redox homeostasis via lipid anabolism^36^, a mechanism induced during cholesterol catabolism^37^. Taken together, these data indicate that in standard media PDIM(+) growth is impaired due to a deficiency of MMCoA. This was supported by measuring the intracellular abundance of MMCoA by LC-MS. In unsupplemented media, MMCoA levels in PDIM(+) *Mtb* were approximately 3-fold lower than in PDIM(-), but increased to similar levels with propionate supplementation (Fig. 3f). Vitamin B_12_ supplementation also significantly increased MMCoA but not propionyl-CoA levels in PDIM(+) *Mtb* (Fig. 3g and Extended Data Fig. 7a), supporting the notion that MMCoA deficiency specifically is responsible for the growth retardation in PDIM(+) *Mtb*.

### Propionate and vitamin B_12_ maintain an intact PDIM biosynthetic pathway

Based on these findings, we reasoned that addition of propionate to culture media would prevent PDIM loss by eliminating the growth advantage of PDIM(-) cells. Whilst arguably cholesterol could also be used for this purpose, propionate is both more affordable and much simpler to work with as a routine media supplement. To assess this, we performed *in vitro* experimental evolution using culture stocks with different PDIM(+) to PDIM(-) ratios. First, we serially passaged H37Rv-B – a moderately PDIM(+) mixed population (Fig. 2f), by weekly subculture in 7H9/OADC/glycerol/tyloxapol ± 0.1 or 1.0 mM propionate and assessed PDIM levels by TLC and VAN10-P assays. Considerable PDIM loss was observed in unsupplemented media whereas propionate supplementation fully maintained PDIM production (Fig. 4a). Repeating this experiment but starting from a PDIM(+) clone, we saw a marked decline in PDIM in unsupplemented media after five passages, whilst 0.1 mM propionate maintained PDIM production for the duration of the experiment (Extended Data Fig. 8c). Strikingly, starting from H37Rv-A – a predominantly PDIM(-) population, PDIM levels progressively increased with propionate (Fig. 4b). VAN10-P screening of single colonies confirmed enrichment of PDIM(+) clones (Fig. 4c), demonstrating 0.1 mM propionate positively selects for PDIM-producing *Mtb*. We speculated that whilst advantageous for the growth of PDIM(+) *Mtb*, 0.1 mM propionate selects against PDIM(-) cells due to propionyl-CoA toxicity^38^ in the absence of a functional PDIM biosynthetic pathway. Indeed, the addition of vitamin B_12_ to alleviate propionyl-CoA toxicity via activation of the methylmalonyl pathway^22,35^ considerably slowed the selection process and resembled more cultures with vitamin B_12_ alone (Fig. 4d and Extended Data Fig. 8d,e). This represents an important advance for the tuberculosis field as it demonstrates that propionate improves the growth of PDIM(+) cells and provides a competitive advantage against spontaneous PDIM(-) mutants, thereby enabling the maintenance of pure PDIM(+) *Mtb* cultures *in vitro*.

**Fig. 4.**
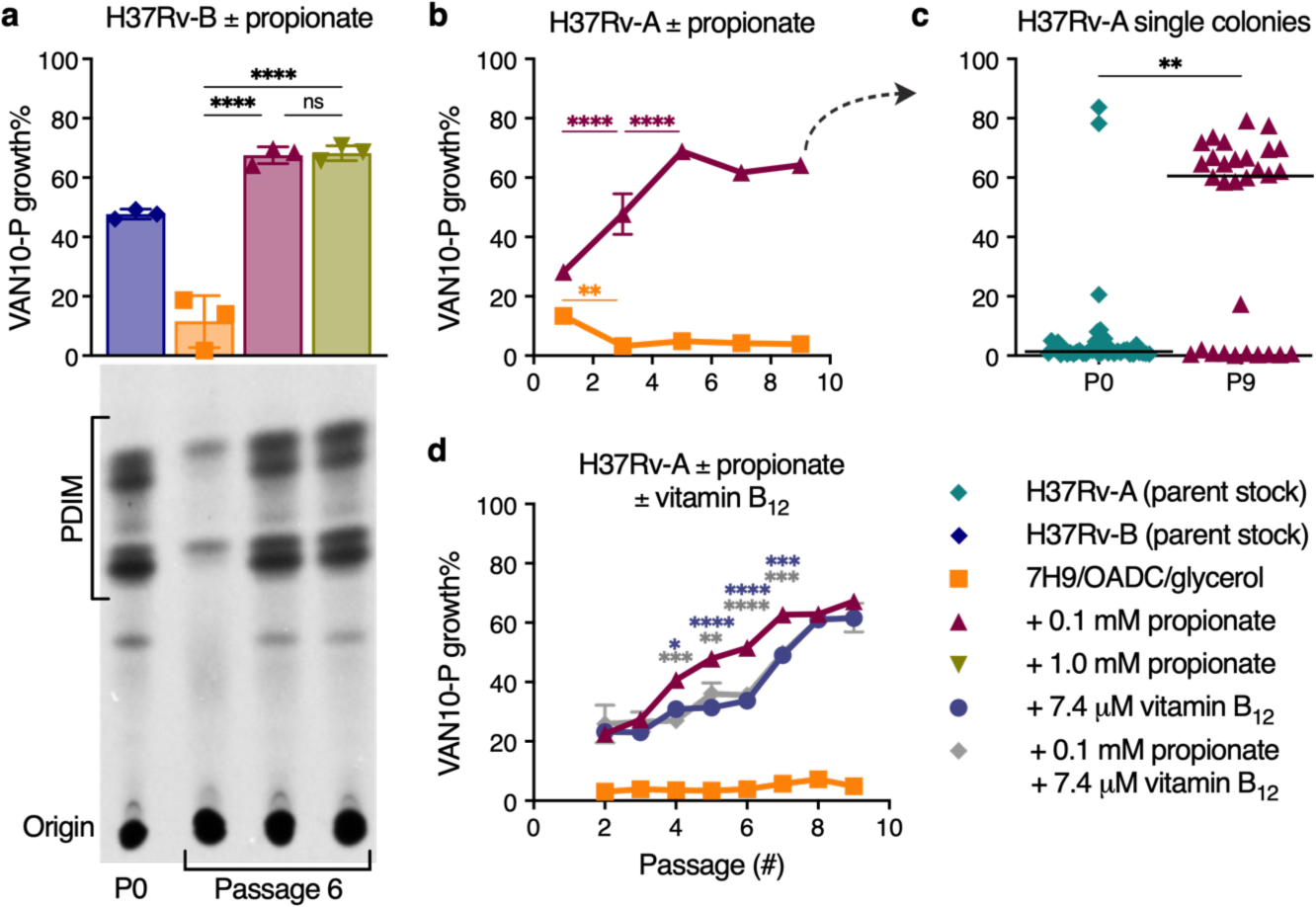
| Propionate and vitamin B_12_ supplementation prevent PDIM loss in *Mtb*. **a**, VAN10-P and TLC lipid analysis of PDIM levels in *Mtb* H37Rv-B following serial passage in 7H9/OADC/glycerol/tyloxapol media ± 0.1 or 1.0 mM propionate. \*\*\*\**P* < 0.0001; one-way ANOVA with Tukey’s multiple comparison test. A representative result is shown for one of two biological replicates analysed by TLC (see also Extended Data Fig. 8b). **b**, VAN10-P assays of H37Rv-A passaged in ± 0.1 mM propionate. **c**, VAN10-P screening of single colonies of H37Rv-A before (*n* = 38; same data as Fig. 2f) and after propionate passage in (**b**) (*n* = 30). Each colony was assayed in triplicate and data points represent mean VAN10-P growth%. Lines indicate the median. *P* = 0.0047; unpaired two-tailed Mann-Whitney test. **d**, VAN10-P assays of H37Rv-A passaged in media supplemented with ± 0.1 mM propionate and 7.4 μM vitamin B_12_ (10 μg/ml) alone and in combination. For (**b**,**d**) \**P* < 0.05, \*\**P* < 0.01, \*\*\**P* < 0.001, \*\*\*\**P* < 0.0001; two- way ANOVA with Tukey’s multiple comparison test. Significant differences are indicated between successive timepoints for each condition in (**b**) and compared to + 0.1 mM propionate in (**d**). *P* > 0.05 for vitamin B_12_ versus vitamin B_12_ + propionate and \*\*\*\**P* < 0.0001 for standard media versus each supplemented condition at all timepoints in (**d**). VAN10-P data in (**a**,**b**,**d**) show mean ± SD for *n* = 3 biological replicates, each assayed in triplicate. For some data points the SD is smaller than the data symbols.

### Propionate increases rifampicin and bedaquiline resistance via enhanced PDIM production

Propionate provides a source of MMCoA precursors that mimics host nutrient conditions but are classically absent in *Mtb* culture media. As PDIM levels had such a profound effect on vancomycin resistance, we sought to further explore the impact of propionate supplementation on drug resistance. MIC assays of several first and second line antitubercular drugs revealed that propionate significantly increased resistance to rifampicin and bedaquiline in a PDIM-dependent manner, while smaller inhibitors like isoniazid, linezolid, and pretomanid showed no difference (Fig. 5 and Supplementary Figs. 5 and 6). Vitamin B_12_ mirrored the effects of propionate supplementation on rifampicin and bedaquiline resistance (Fig. 5a,c), consistent with enhanced resistance due to increased PDIM production. Notably, the rifampicin MIC_90_ for PDIM(+) *Mtb* in propionate-supplemented media reduced ∼30-fold when tyloxapol was replaced with Tween 80 (Fig. 6), arguing that Tween 80 is unsuitable for detecting *in vivo* relevant drug sensitivity. These data are also congruent with our earlier results showing greater resistance of PDIM(+) *Mtb* to large compounds and lower permeability compared to PDIM(-) (Extended Data Fig. 1). One outlier to this trend was capreomycin (668.7 g/mol), which did not exhibit a PDIM-propionate MIC shift (Supplementary Fig. 5a). Wang *et al*. have also reported PDIM-dependent resistance to the small molecule inhibitor 3bMP1 (229.3 g/mol)^12^, indicating that the relationship between the PDIM permeability barrier and drug uptake is complex though the uptake of large compounds is more likely to be affected by PDIM.

**Fig. 5.**
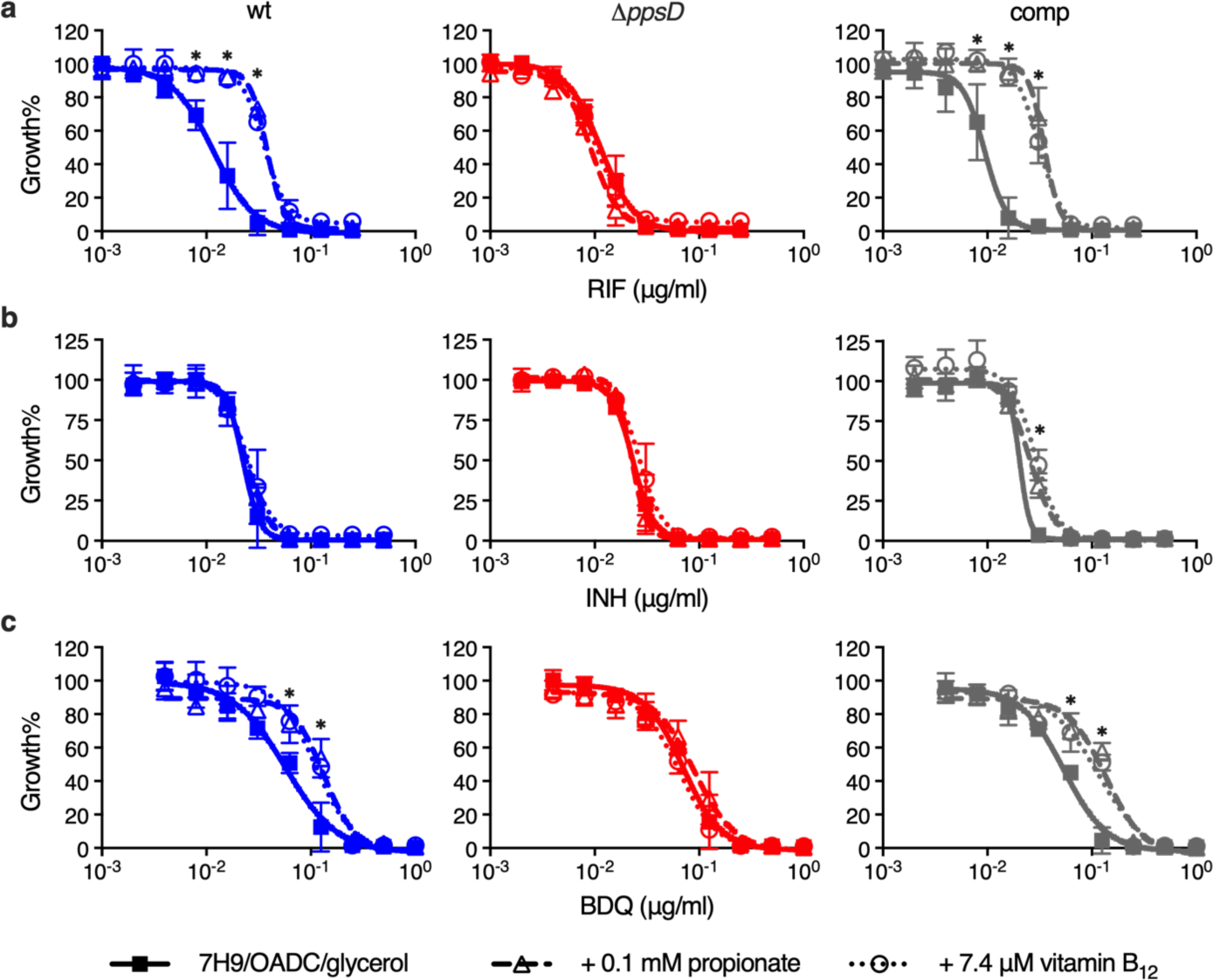
| Propionate and vitamin B_12_ supplementation increase rifampicin and bedaquiline resistance of *Mtb* in a PDIM-dependent manner. **a**, Sensitivity of PDIM(+) and PDIM(-) *Mtb* H37Rv to rifampicin (RIF), **b**, bedaquiline (BDQ), and **c**, isoniazid (INH), in standard 7H9/OADC/glycerol/tyloxapol media and supplemented with either 0.1 mM propionate or 7.4 μM vitamin B_12_ (10 μg/ml). \**P* < 0.001 for both propionate and vitamin B_12_ versus unsupplemented; two-way ANOVA with Tukey’s multiple comparison test. Mean ± SD for *n* = 4 biological replicates from two independent experiments.

**Fig. 6.**
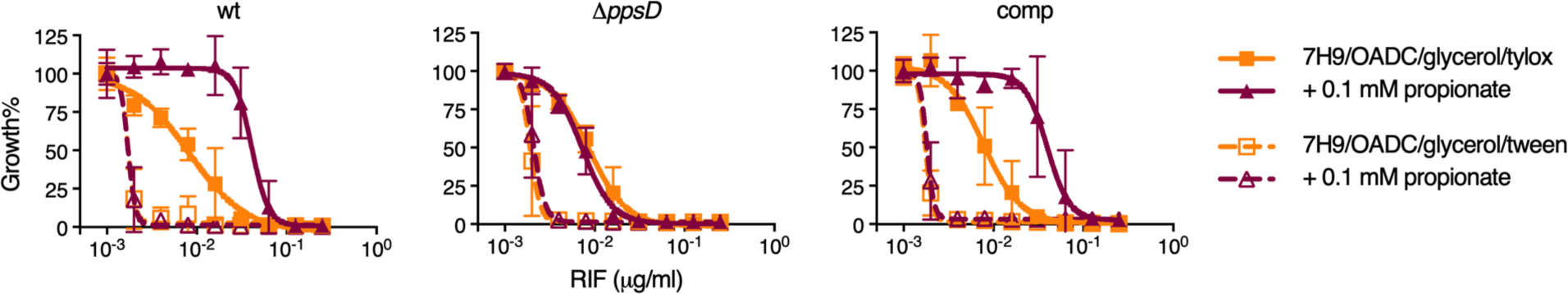
| Tween 80 increases the sensitivity of *Mtb* to rifampicin and abolishes PDIM- dependent differences in MIC. Sensitivity of PDIM(+) and PDIM(-) *Mtb* H37Rv to rifampicin (RIF) in 7H9/OADC/glycerol ± 0.1 mM propionate using either tyloxapol or Tween 80 as the culture detergent. Mean ± SD for *n* = 4 biological replicates from two independent experiments.

## Discussion

The mycobacterial cell wall plays a crucial role in the interactions between the pathogen and host^39^. However, the study of *Mtb* cell wall biology and pathogenesis has long been impeded by PDIM bias. It has been nearly 50 years since the first report associating PDIM loss with virulence attenuation *in vivo*^8^ and over 20 years since PDIM loss and attenuation were linked at the genetic level^6,7^. Numerous studies have since been published reporting spontaneous PDIM loss not only in *Mtb* H37Rv^3–5,12^, but also in *Mtb* Erdman^40^, HN878^3^, CDC1551^29,41^, and in *M. bovis* BCG vaccine strains^42^. The PDIM problem is thus both long-standing and far-reaching, and the number of studies unpublished due to PDIM bias and the time and resources spent chasing PDIM-related phenotypes are high.

We discovered that the cause and solution of the PDIM problem are rooted in *Mtb* metabolism. *Mtb* growth is impaired in the absence of an exogenous source of MMCoA precursors, providing a selection pressure for PDIM loss. This bottleneck can be alleviated by supplementing propionyl-CoA-generating carbon sources such as odd-chain fatty acids or cholesterol, or the cofactor vitamin B_12,_ which increase MMCoA pools and restore full growth of PDIM(+) *Mtb*. The affinity of *Mtb* for host fatty acids was first described by Segal and Bloch in 1956^43^ and today we know that *Mtb* is a specialist in cholesterol utilization^18^. Cholesterol is thought to be a major source of propionyl-CoA in the host^19^, and *Mtb* has evolved to efficiently use this host-derived resource as fuel for cellular metabolism and a building block for virulence lipids^18,35^. Moreover, starving the bacterium of propionyl-CoA via cholesterol limitation improves macrophage control of *Mtb*^44^, demonstrating a crucial role for this metabolite in *Mtb* during infection. Why then, to this day, has *Mtb* culture media remained devoid of propionyl-CoA precursors? Most *Mtb* culture media were optimized to promote rapid planktonic growth rather than to reflect the nutrient environment found in the host^45,46^, and we speculate the toxicity of high concentrations of odd-chain fatty acids has previously discouraged their inclusion. Contrary to its negative reputation, we show that 0.1 mM propionate is in fact advantageous for PDIM(+) growth and selects against PDIM(-) cells analogous to animal passage, providing an elegant and long-sought solution to the PDIM problem. Our data also clearly demonstrate that tyloxapol and not Tween 80 should be used in PDIM assays and PDIM selective media as tyloxapol maintains PDIM-dependent impermeability, whilst Tween 80 strips PDIM and other cell wall components^27^.

These data also have ramifications for future drug discovery efforts. During host infection, *Mtb* survives in a PDIM-rich state^17^, however, PDIM production is poorly supported in current culture media potentially leading to overestimation of drug potency. Indeed, we show that PDIM levels affect the potency of rifampicin, bedaquiline and other high molecular weight inhibitors. In concert with the current literature^47,48^, this clearly points to the mycobacterial cell wall as an important factor in drug efficacy. In addition, the tremendous increase in vancomycin and rifampicin sensitivity with Tween 80, together with previous reports of this phenomenon^49,50^, strongly advocate for the use of tyloxapol to maintain the natural permeability barrier in *Mtb*. Our findings also expand on previous observations associating propionyl-CoA metabolism with rifampicin resistance^25,47^ by directly linking propionyl-CoA and PDIM production with enhanced rifampicin resistance. The decreased virulence and increased drug sensitivity of PDIM(-) *Mtb* suggest that inhibitors of PDIM biosynthesis could significantly increase the *in vivo* potency of current drug regimens. Such a therapeutic option would be highly specific, as PDIMs are confined to slow-growing, pathogenic mycobacteria. Furthermore, inhibitors targeting propionyl-CoA metabolism could be synergistic with rifampicin due to downstream effects on PDIM.

The main recommendations stemming from our study are to routinely supplement culture media with 0.1 mM propionate and avoid Tween 80 to prevent PDIM loss and the emergence of heterogeneous populations whilst simultaneously augmenting PDIM production. VAN-P assays provide the tools for routine strain testing during genetic manipulations and enable efficient re-isolation of PDIM(+) strains from mixed populations. Pure and properly maintained PDIM(+) strains will be indispensable for studying interactions with host immunity, as well as for high-stakes pre-clinical work such as vaccine studies in non-human primates. Moreover, today in the emerging era of *Mtb* systems biology where large pools of genetically modified strains are crucial tools for *in vitro* and *in vivo* studies^51–53^, preventing secondary PDIM mutations is imperative. Importantly, our approach is accessible to everyone, including labs in low-resource settings.

Taken together, our discoveries not only solve the PDIM problem, but also highlight how the host nutritional environment has shaped the close coupling between *Mtb* metabolism and its virulence. Our findings exemplify how discrepancies between the host and *in vitro* nutrient environment can attenuate bacterial pathogenicity and provide essential tools and culture conditions to finally eliminate the PDIM problem from tuberculosis research.

## Methods

### Bacterial strains, culture condition and reagents

*Mtb* strains were obtained from laboratory stocks and are listed in Supplementary Tables 1 and 2. Fresh starter cultures were inoculated from frozen seed stocks and then subcultured once before use in experiments. Subcultures were typically grown for four days to an optical density (OD) at 600 nm (OD_600_) of ∼0.8. For BSL2 strains, OD_600_ was measured using a GENESYS 140 spectrophotometer (Thermo Fisher Scientific). For BSL3 strains, OD_600_ was measured on a Biowave WPA CO8000 spectrophotometer (Biochrom Ltd.) and then converted using a calibration curve constructed against a GENESYS 10uv spectrophotometer (Thermo Fisher Scientific). Preculturing steps were performed using Middlebrook 7H9 broth supplemented with 10% (v/v) OADC (0.6 g/l sodium oleate, 50 g/l bovine serum albumin fraction V, 20 g/l dextrose, 40 mg/l catalase, 8.5 g/l sodium chloride), 0.2% (v/v) glycerol, and 0.05% (v/v) tyloxapol. This is referred to as standard 7H9/OADC/glycerol/tyloxapol media. BSL2 strains (Supplementary Table 1) were additionally supplemented with 24 mg/l D-calcium pantothenate, 200 mg/l L-arginine, 50 mg/l L-leucine, and 50 mg/l L-methionine (‘PALM’ supplements). Hygromycin B at 75 μg/ml and kanamycin at 30 μg/ml were added to precultures as indicated (Supplementary Table 2). For supplemented media, 1000 × supplement stocks were prepared in MilliQ water, filter sterilized, then added to standard media and the pH checked. Final supplement concentrations were as follows: 0.1 mM or 1.0 mM sodium propionate, 7.4 μM vitamin B_12_ (10 μg/ml), and 0.1 mM sodium pyruvate, sodium acetate, sodium butyrate and valeric acid. Cholesterol was prepared at 0.1 M in 1:1 (v/v) EtOH/tyloxapol as previously described^35^ and then added to detergent-free media to give a final concentration of 0.1 mM cholesterol and 0.05% tyloxapol. Controls were prepared by adding EtOH/tyloxapol in the same manner to provide the detergent. For Tween 80 experiments, 0.05% Tween 80 was used in place of tyloxapol. For growth curve experiments, triplicate inkwells with 5 ml of media were inoculated at a starting OD_600_ of 0.01. Broth cultures were grown at 37 °C with gentle shaking (100 rpm for BSL2 strains, 80 rpm for BSL3). Middlebrook 7H10 agar supplemented with 10% (v/v) OADC and 0.5% (v/v) glycerol (7H10/OADC/glycerol) was used as a solid media for plating and plates were incubated at 37 °C for three weeks. Supplier information for media components and supplements are listed in Supplementary Table 3.

### Mutant generation and complementation

Deletion of the *tgs1* (*Rv3130c*), *ppsD* (*Rv2934*) and *mas* (*Rv2940c*) genes was carried out by specialized transduction as previously described^28^. H37Rv-B was used to generate H37Rv 11.*tgs1* mutants; H37Rv-SC [a single PDIM(+) clone isolated from H37Rv-B by VAN10-P screening] was used to generate H37Rv 11.*ppsD*; and CDC1551-B to generate CDC1551 11.*mas* (Supplementary Table 2). Transductants were selected on plates containing hygromycin (75 μg/ml) and the deletion was confirmed by 3-primer PCR and whole genome sequencing (WGS). The 11.*ppsD* strain was complemented using the integrative vector pMV361^54^ containing a copy of the *ppsD* gene under control of the HSP60 promoter (pMV361-*ppsD*). The complementation plasmid was constructed by Gibson assembly using the NEBuilder HiFi DNA Assembly Cloning Kit (New England Biolabs). In brief, the plasmid and *ppsD* insert were amplified by PCR and a Gibson assembly reaction was used to transform *Escherichia coli* DH5α. The plasmid was isolated, and the nucleotide sequence of the construct was verified by Sanger sequencing. H37Rv 11.*ppsD* cells were electroporated with ∼0.5 μg of the complementation plasmid, recovered overnight in 7H9/OADC/glycerol/tyloxapol at 37 °C with shaking and then selected on plates containing hygromycin (75 μg/ml) and kanamycin (30 μg/ml). The H37Rv 11.*ppsD*::comp strain was validated by PCR to confirm both the complementation and presence of the 11.*ppsD* deletion. Primers used for vector construction and PCR confirmation are listed in Supplementary Table 4.

### Thin layer chromatography

*Mtb* cultures were grown to early log phase and then diluted to OD_600_ 0.3 in 10 ml 7H9/OADC/glycerol/tyloxapol and labelled with propionic acid [1-^14^C] sodium salt (7 μCi) (American Radiolabeled Chemicals, Inc.). Cultures were incubated with shaking at 37 °C for two days and then spun down. Methanol (2.0 ml), 0.3% sodium chloride aqueous solution (0.2 ml) and petroleum ether (2.0 ml) were added to the cell pellets and the suspensions were vortexed for 30 s followed by centrifugation. The petroleum ether phases were moved to new tubes and the extraction with petroleum ether was repeated twice. The petroleum phases were combined, dried with anhydrous sodium sulfate, filtered and evaporated to dryness under nitrogen. The PDIM extracts were resuspended in dichloromethane (0.2 ml). Counts per minute (cpm) were measured to load approximately 5000 cpm for each sample on a silica gel 60 F254 thin layer chromatography (TLC) plate (Sigma-Aldrich). The TLC plate was eluted three times with petroleum ether/ethyl acetate 98/2. PDIMs were detected by autoradiograph after exposure for 48–72h at −80 °C. PDIM band intensity was quantified using ImageJ (v 1.52a)^55^.

### MIC assays

Resistance of *Mtb* strains to vancomycin and other inhibitors (Supplementary Table 5) were determined using the microbroth dilution method. Two-fold serial dilutions at 2 × final drug concentration were prepared in standard 7H9/OADC/glycerol/tyloxapol or media supplemented with either 2 × propionate (0.2 or 2.0 mM) or vitamin B_12_ (14.8 μM) at a volume of 100 μl in the inner wells of flat-bottom 96-well plates. The outer wells were aliquoted with 200 μl PBS or media. Strains were precultured in 7H9/OADC/glycerol/tyloxapol to OD_600_ of ∼0.8 and then diluted to OD_600_ 0.01 in the same media. 100 μl of the cell dilution was added to plate to give a final OD_600_ of 0.005; 0.1 or 1.0 mM propionate or 7.4 μM vitamin B_12_ for supplemented assays; and 1 × drug concentration. Plates were incubated with gentle shaking and bacterial growth was measured by OD after 10 days unless otherwise specified. For BSL2 strains, OD_600_ was measured on a FLUOstar Omega Microplate Reader (BMG LABTECH). For BSL3 strains, OD_590_ was measured on an Epoch BioTek Microplate Spectrophotometer (BioTek Instruments, Inc.). Data were normalized to drug-free control wells and fit with non-linear regression in Prism (v9.4.1, v10.0.1) (GraphPad Software). MIC_90_ and MIC_50_ values were calculated from the curve fit.

### VAN10 assay

VAN10 assays were performed in the inner wells of flat-bottom 96-well plates prepared with standard 7H9/OADC/glycerol/tyloxapol or media supplemented with 2 × propionate (0.2 or 2.0 mM) or vitamin B_12_ (14.8 μM). Triplicate wells were aliquoted with 100 μl drug-free media or media with 20 μg/ml vancomycin. Strains were precultured as for MIC assays and diluted to an OD_600_ of 0.01 in 7H9/OADC/glycerol/tyloxapol. 100 μl of the cell dilution was added to the plate giving a final vancomycin concentration of 10 μg/ml in treated wells (VAN10); OD_600_ of 0.005; and 0.1 or 1.0 mM propionate or 7.4 μM vitamin B_12_ in supplemented assays. Plates were incubated with gentle shaking and bacterial growth was measured by OD after 10 days unless otherwise specified. Relative growth in VAN10 was calculated compared to drug-free wells (VAN0) (VAN10 OD / VAN0 OD × 100 = VAN10 growth%). The VAN10 assay supplemented with 0.1 mM propionate is referred to as the ‘VAN10-P’ assay.

For high throughput screening of single colonies and to isolate PDIM(+) clones, single colonies were picked into 7H9/OADC/glycerol/tyloxapol and grown until dense to synchronize. Outgrowth cultures were then subcultured for a single passage and grown to an OD_600_ of ∼0.5–1.0. Subcultures were diluted 1:50 in 7H9/OADC/glycerol/tyloxapol and 100 μl of this was used to inoculate VAN10-P assay plates prepared as above. Growth was measured after 14 days to obtain an endpoint measurement. mc^2^6230 was additionally supplemented with 24 mg/l pantothenate, and 0.1 mM propionate was included in the plates and outgrowth media used to isolate mc^2^6230 AE1601 (Supplementary Table 1).

### Permeability assay

Cell envelope permeability was determined using the Ethidium Bromide (EtBr) uptake assay^16^. Four replicate cultures of *Mtb* mc^2^7902 and mc^2^8398 in 10 ml 7H9/OADC/glycerol/ tyloxapol + PALM media were grown to an OD_600_ of 0.6–1.0. Cultures were washed three times with PBS + 0.4% (w/v) glucose and diluted to an OD_600_ of 0.5. Five replicate 180 µl aliquots were transferred to a black, clear-bottom, 96-well plate and 20 µl EtBr (50 μg/ml) was added. The plate was incubated at 37 °C in a FLUOstar Omega Microplate Reader (BMG LABTECH) with 300 rpm double-orbital shaking. Fluorescence was measured at an excitation wavelength of 355 nm and emission wavelength of 590 nm every 15 min for one hour.

### Evolution experiments

Triplicate inkwells containing 10 ml standard 7H9/OADC/glycerol/tyloxapol or supplemented media as specified were inoculated with 100 μl of frozen *Mtb* seed stock and incubated for 7–10 days. Cultures were then diluted 1:250 into 10 ml fresh media each week for serial passage. To assess PDIM maintenance over the course of the experiment, at selected passages cultures were input into VAN10-P assays and 1 ml of culture was stocked and stored at −80 °C. VAN10-P assay plates were prepared as above and cultures were diluted 1:100 in 7H9/OADC/glycerol/tyloxapol for input into the assay. Growth was measured after 7 and 14 days of incubation. For TLC lipid analysis of passaged cultures, cultures were first recovered from frozen stocks by growing to an OD_600_ of ∼1.0 in standard 7H9/OADC/glycerol/tyloxapol before ^14^C-labelling and TLC lipid analysis as above.

### Metabolomics extractions

Triplicate inkwells containing 7 ml standard 7H9/OADC/glycerol/tyloxapol or supplemented media as specified were inoculated at OD_600_ 0.01 and grown for five days and then harvested. An equivalent of 3 ml culture at an OD_600_ of 1.0 was rapidly filtered on 0.45 μm Durapore PVDF membrane filters (MilliporeSigma) using a vacuum manifold (MilliporeSigma). Cultures were quenched by placing the filter paper in 1 ml of extraction solvent containing 20:40:40 (v/v) water/acetonitrile/methanol with approximately 500 μl of 0.1 mm zirconia/silica beads (BioSpec) at −20 °C. Samples were homogenized using a Precellys Cryolys Evolution (Bertin Technologies) cooled to 0 °C for three 20 s cycles at 6800 rpm with a 30 s pause between cycles. Samples were centrifuged and the extracts were filtered through a 0.22 μm Nylon Spin-X microcentrifuge filter (Corning) and stored at −80 °C. For analysis, extract samples were concentrated 5-fold by using a SpeedVac^®^ Plus SC110A (Savant Instruments, Inc.) to evaporate the solvent and then redissolved in 1/5^th^ volume of the extraction solvent.

### LC-MS metabolomic profiling

Metabolomics analysis was performed using an Agilent 1290 Infinity II liquid chromatography (LC) system coupled with an Agilent 6545 quadrupole time-of-flight (QTOF) mass spectrometer (MS) equipped with a Dual Agilent Jet Stream Electrospray Ionization (Dual AJS ESI) source operated in negative mode. Metabolites were separated on an InfinityLab Poroshell 120 HILIC-Z, 2.1 x 150 mm, 2.7 µm, 100 Å column (Agilent) based on previously described methods^56^. The mobile phase consisted of solvent A: water, and solvent B: 15:85 (v/v) water/acetonitrile, both with 10 mM ammonium acetate and 2.5 μM InfinityLab Deactivator Additive (Agilent), pH 9.0. HPLC grade water (Cen-Med Enterprises) and LC-MS grade solvents (Fisher Chemical) were used for both the LC-MS mobile phase and metabolite extraction. The elution gradient used was as follows: 0–2 min 96% B; 2–5.5 min 96 to 88% B; 5.5–8.5 88% B; 8.5–9 min 88 to 86% B; 9–14 min 86% B; 14–17 min 86 to 82% B; 17–23 min 82 to 65% B; 23– 24 min 65% B; 24–24.5 min 64 to 96% B; 24.5–26 min 96% B; followed by a 3 min re-equilibration at 96% B. The flow rate was 0.25 ml/min and column temperature 50 °C. The injection volume was 3 μl and the autosampler was maintained at 4 °C during the run. Mass spectra were recorded in profile mode from *m*/*z* 60 to 1200 using an acquisition rate of 1 spectra/s in the 2GHz extended dynamic range mode and 1700 m/z low mass range, using the sensitive slicer mode and fragile ions option. The gas temperature was 225 °C and sheath gas temperature 350 °C. The capillary, nozzle, fragmentor, skimmer, and octopole voltages were 3500, 2000, 125, 45 and 750 V, respectively. Dynamic mass axis calibration was achieved by continuous infusion of a reference mass solution using an isocratic pump with a 100:1 splitter.

Data Analysis was performed using the Agilent MassHunter Qualitative and Quantitative Analysis Software. Metabolite identification was based on mass-retention times determined using chemical standards (Supplementary Table 6) and isotope distribution patterns. Calibration curves of standard compound mixtures in extraction buffer and spiked into a homologous mycobacterial extract were run to determine the linear range. Metabolites were quantified using a mass tolerance of 20 ppm with manual curation of peak areas where necessary and the area under the curve (AUC) was determined. AUC was normalized using the median total AUC for a panel of 50 putative metabolites across different metabolic pathways (Supplementary Table 7) to correct for differences in extraction and concentration efficiency.

### Mouse experiments

Mouse experiments were performed in accordance with National Institutes of Health guidelines following the recommendations in the Guide for the Care and Use of Laboratory Animals^57^. The protocols used in this study were approved by the Institutional Animal Care and Use Committee of Albert Einstein College of Medicine (Protocols #00001445 and #00001332). To generate H37Rv-B, female C57BL/6 mice (Jackson Laboratory) were infected with H37Rv-A via the aerosol route using a 1 × 10^7^ cfu/ml *Mtb* suspension in PBS containing 0.05% tyloxapol and 0.004% antifoam. Mice were sacrificed after 21 days and the lungs homogenized and plated on 7H10/OADC/glycerol plates. All colonies from the lung of a single mouse were harvested and used to inoculate 7H9/OADC/glycerol/tyloxapol in a roller bottle. This was grown to an OD_600_ of 1.8 and then stocked in 1 ml aliquots and stored at −80 °C. To isolate single PDIM(+) clones of Erdman, HN878 and CDC1551, in-house bred Rag^-/-^ mice were infected with 5 × 10^6^ cfu/mouse via the intravenous route. Mice were killed on day 20 post-infection and the lungs homogenized and plated on 7H10/OADC/glycerol plates. Single colonies were picked and outgrown in 7H9/OADC/glycerol/tyloxapol with 0.1 mM propionate for stocking and then subcultured and screened for PDIM using VAN10-P assays as above.

### Whole genome sequencing and analysis

Genomic DNA was isolated using a CTAB extraction method as previously described^58^ and sequenced in-house on an Illumina MiSeq. Genomic libraries were prepared using the Illumina Nextera XT library preparation kit and sequenced with a 600-cycle v3 reagent kit (2 × 301 bp reads) following the manufacturer’s instructions. Genomes with uneven coverage (< 90% of the genome having > 10 × coverage) for which no PDIM SNPs were detected were additionally sequenced with a 150-cycle v3 kit (2 × 76 bp reads) and the data merged for mapping. Additional sequencing by Illumina NextSeq was performed by SeqCenter (Pittsburgh, PA) using the Illumina DNA Prep kit and sequenced on an Illumina NextSeq 2000 (2 × 151 bp reads).

Raw reads were trimmed with Trimmomatic (v0.39)^59^ using a sliding window quality filter (SLIDINGWINDOW:4:15) and reads less than 25 bp were discarded (MINLEN:25). Trimmed reads were then mapped to the reference genome corresponding to the strain background (H37Rv NC_000962.3, CDC1551 NC_002755.2, Erdman NC_020559.1, HN878 NZ_CM001043.1 and KZN 4207 NC_016768.1) using BWA-MEM (v0.7.17-r1188) (https://github.com/lh3/bwa). Mapping files were sorted and indexed using Samtools (v1.6)^60^. Duplicates were removed using Picard tools (v2.26.10) (http://broadinstitute.github.io/picard) and local realignment was performed using GATK (v.3.8-0)^61^. Mapping quality was assessed using Qualimap (v2.2.1)^62^. Variants were called using Pilon (v1.23)^63^ using a minimum depth threshold of 5, base quality threshold of 15 and mapping quality threshold of 40 (--variant --mindepth 5 --minqual 15 --minmq 40). Variants were annotated using SNPeff (v5.1d)^64^. Geneious Prime® (v2022.2.2) (Biomatters Ltd.) was used to detect low-frequency variants within the PDIM gene region (*tesA–Rv2953*) using the variation/SNP finder feature with a coverage threshold of 10, minimum variant frequency of 10%, and *P* value < 1 x 10^-10^. SeqTK (v1.3-r106) (https://github.com/lh3/seqtk) was used to randomly subsample reads for downsampling analyses.

### *ppsC* homopolymeric tract region Sanger sequencing

To identify and confirm mutations in the *ppsC* homopolymeric tract region, a 250 bp fragment encompassing this region was amplified by PCR and then sequenced by Sanger sequencing. PCR was performed in 50 μl reactions containing 2.5 units HOT FIREPol^®^ DNA polymerase (Solis BioDyne), the supplied reaction buffer BD at 1 × concentration, 2.0 mM MgCl_2_, 250 μM dNTPs, 0.3 μM of each primer, and 2.5% (v/v) DMSO. Primers are listed in Supplementary Table 4. Approximately 25 ng of gDNA was used as the PCR template. Thermal cycling consisted of an initial denaturation and enzyme activation step of 15 min at 95 °C, followed by 35 cycles of 30 s at 95 °C, 45 s at 55 °C, and 30 s at 72 °C. This was followed by a final elongation step of 10 min at 72 °C. PCR products were purified using the Wizard® SV Gel and PCR Clean-Up system (Promega) and then sequenced by Sanger sequencing at GENEWIZ (South Plainfield, NJ) in both the forward and reverse direction using the same primers as for amplification.

### Statistical analysis

Statistical analyses were performed using Prism (v9.4.1 and v10.0.) (GraphPad Software). Significant differences were calculated by one- or two-way ANOVA using multiple comparison tests as specified, or the nonparametric Mann-Whitney test for skewed data. Correlations between vancomycin MIC and VAN10 growth% with PDIM were assessed by simple linear regression.

### Data availability

Whole genome sequence data have been deposited in the NCBI Sequence Read Archive (SRA) under the BioProject accession number PRJNA923717. A complete list of strains sequenced in this study and SRA accession numbers are given in Supplementary Table 8. Raw metabolomics data are provided as a source data file.

## Acknowledgments

We thank Bing Chen, John Kim, and Mei Chen for assistance with animal experiments; Annie Zhi Dai for technical support; and the labs of Jeremy Rock, Rockefeller University, NY, and John Chan, Rutgers University, NJ, for their feedback and independent validation of VAN-P PDIM assays. C.V.M., T.J.W., J.C., E.Z.R., and M.B. acknowledge support from the National Institutes of Health/National Institute of Allergy and Infectious Diseases (R01 AI139465 and R01 AI175972), the Potts Memorial Foundation, and Albert Einstein College of Medicine internal funding. M.W.S. acknowledges support from the Institutional AIDS training grant, Training in HIV/AIDS Pathogenesis; Basic and Translational Research (T32 AI007501), and the Albert Einstein College of Medicine MSTP training grant.

## Author contributions

C.V.M. and M.B. conceived and designed the study. C.V.M., T.J.W., J.C., C.V., S.R., M.W.S., and E.Z.R., performed the experiments. C.V.M., T.J.W., C.V., M.W.S., and M.B. analysed the data. M.B. and W.J.R. provided resources. C.V.M. and M.B. wrote the paper. T.J.W., C.V., S.R., M.W.S., E.Z.R., and W.J.R. critically reviewed and edited the paper.

## Additional information

Supplementary Information is available for this paper.

Correspondence and requests for materials should be addressed to M.B.

Reprints and permissions information is available at www.nature.com/reprints

---

**Extended Data Table 1.** | PDIM mutations in *Mtb* strains included in this study. Non-synonymous variants in genes involved in PDIM biosynthesis, assembly, processing, or transport (see Fig. 2e). Mutations are described at the protein level for amino acid substitutions (p.) and at the gene coding level for frameshift/INDEL mutations (c.). Variants were identified from Illumina MiSeq WGS data using the software tool Pilon^63^ for variant calling unless indicated. For mutations with mixed coverage (< 90%) the frequency of the variant allele is given in brackets. ‘H37Rv-B col#’ are single colonies isolated from H37Rv-B with different VAN10-P growth%, including a subset of those which had an intermediate VAN10-P phenotype (see Fig. 2f). ‘≥ 10 ξ’ is the percentage of genome with 10 ξ or greater coverage.

| Strain | Genome coverage (x) |  |  | PDIM mutations |
| --- | --- | --- | --- | --- |
|  | Mean | SD | ≥ 10 x |  |
| mc<sup>2</sup>6206 | 66.3 | 17.4 | 99.00% | n.d. |
| mc<sup>2</sup>6209 | 56.8 | 21.8 | 98.90% | n.d. |
| mc<sup>2</sup>6230 | 187.0 | 73.8 | 98.50% | *ppsC* c.2685(C)<sub>7→8</sub> (66%)† |
| mc<sup>2</sup>6230 AE1601* | 171.7 | 32.6 | 99.11% | n.d. |
| mc<sup>2</sup>6230 AE1611 | 30.5 | 9.4 | 97.26% | *ppsC* c.2685(C)<sub>7→8</sub> ‡ |
| mc<sup>2</sup>7901 | 26.7 | 10.5 | 96.50% | *fadD28* p.Arg562Gly |
| mc<sup>2</sup>7902 | 47.8 | 15.9 | 98.50% | n.d. |
| mc<sup>2</sup>8398 | 62.1 | 16.7 | 99.20% | *ppsC* c.2475delC |
| H37Rv-A | 93.5 | 31.0 | 97.60% | n.d. |
| H37Rv-B | 60.5 | 18.3 | 97.80% | n.d. |
| H37Rv-SC (AE1028)* | 57.1 | 19.6 | 98.90% | n.d. |
| H37Rv Δ*ppsD* | 58.7 | 18.6 | 98.70% | Δ*ppsD* |
| H37Rv Δ*tgs1-1* | 95.6 | 56.5 | 91.70% | *fadD28* p.Leu241Pro |
| H37Rv Δ*tgs1-2* | 90.3 | 78.6 | 83.00% | *ppsA* p.Trp968* |
| H37Rv Δ*tgs1-3* | 17.2 | 7.0 | 87.20% | *ppsD* p.Ser247Pro |
| H37Rv Δ*tgs1-4* | 86.3 | 25.1 | 97.70% | *ppsE* p.Asp74Asn |
| H37Rv Δ*tgs1-5* | 119.4 | 57.2 | 96.50% | *ppsC* c.2668(C)<sub>7→6</sub> ‡ |
| H37Rv Δ*tgs1-7* | 184.5 | 138.4 | 93.10% | n.d. |
| H37Rv Δ*tgs1-8* | 58.2 | 15.1 | 99.00% | *ppsC* c.2668(C)<sub>7→8</sub> ‡ |
| H37Rv Δ*tgs1-9* | 69.7 | 18.0 | 99.10% | *ppsC* c.2668(C)<sub>7→6</sub> ‡ |
| CDC1551-A | 106.5 | 49.8 | 96.90% | n.d. |
| CDC1551-B | 54.7 | 19.6 | 98.80% | n.d. |
| CDC1551 Δ*mas* | 44.0 | 18.6 | 96.50% | Δ*mas* |
| Erdman | 84.7 | 25.2 | 98.00% | *ppsA* c.3418insA (72%) |
| HN878 | 76.1 | 44.3 | 93.10% | n.d. |
| KZN 4207 | 86.7 | 38.1 | 97.10% | n.d. |
| CDC1551-SC (AE5005)* | 69.3 | 18.9 | 99.10% | n.d. |
| Erdman-SC (AE3003)* | 91.2 | 23.5 | 99.30% | n.d. |
| HN878-SC (AE8005)* | 65.9 | 24.4 | 97.90% | n.d. |
| H37Rv-B col5 (VAN10-P 41%) | 61.0 | 15.7 | 99.00% | *ppsE* p.Asp74Asn |
| H37Rv-B col6 (VAN10-P 1%) | 53.9 | 14.8 | 98.70% | *ppsA* p.Trp968* |
| H37Rv-B col16 (VAN10-P 43%) | 50.7 | 13.7 | 98.60% | *ppsE* p.Asp74Asn |
| H37Rv-B col17 (VAN10-P 77%) | 34.9 | 13.0 | 97.90% | n.d. |
| H37Rv-B col19 (VAN10-P 31%) | 94.0 | 25.0 | 99.10% | *ppsE* p.Asp74Asn |
| H37Rv-B col23 (VAN10-P 39%) | 38.2 | 11.2 | 98.10% | *ppsE* p.Asp74Asn |
| H37Rv-B col36 (VAN10-P 1%) | 62.9 | 15.7 | 98.90% | *ppsA* p.Trp968* |

\*PDIM(+) clone identified by VAN10-P screening of single colonies (see also Extended data Fig. 4). †Mutation identified using the Illumina NextSeq platform; ‡Mutation detected/confirmed by PCR and Sanger sequencing (see also Extended data Fig. 5). n.d. = no non-synonymous PDIM mutations detected.

**Extended Data Table 2.** | Low-frequency PDIM mutation analysis. Low-frequency non-synonymous PDIM variants detected in laboratory stocks of virulent *Mtb* strains (Fig. 2a). Variant calling was performed using the Geneious variant finder with the following thresholds: ≥ 10% variant frequency, ≥ 10 ξ coverage, *P* < 1 x 10^-10^. n.d. = none detected.

| Strain | PDIM mutations | Frequency | REF/ALT | P value |
| --- | --- | --- | --- | --- |
| H37Rv-A | *ppsB* p.Leu250Pro | 11.6% | 99/13 | $3.1 \times 10^{-23}$ |
| H37Rv-B | n.d. |  |  |  |
| Erdman | *ppsA* c.3418insA | 67.9% | 27/57 | $2.9 \times 10^{-161}$ |
| | *ppsE* p.Cys414* | 10.7% | 50/6 | $3.2 \times 10^{-14}$ |
| KZN 4207 | n.d. |  |  |  |
| HN878 | *fadD28* c.347delT | 14.3% | 126/21 | $2.6 \times 10^{-30}$ |
| | *ppsD* p.Gly1636Arg | 18.8% | 26/6 | $1.4 \times 10^{-14}$ |
| CDC1551-A | *ppsA* p.Phe327Val | 10.2% | 115/13 | $7.1 \times 10^{-20}$ |
| | *ppsE* c.3496insA | 12.6% | 118/17 | $9.9 \times 10^{-41}$ |
| CDC1551-B | n.d. |  |  |  |

**Extended Data Fig. 1.**
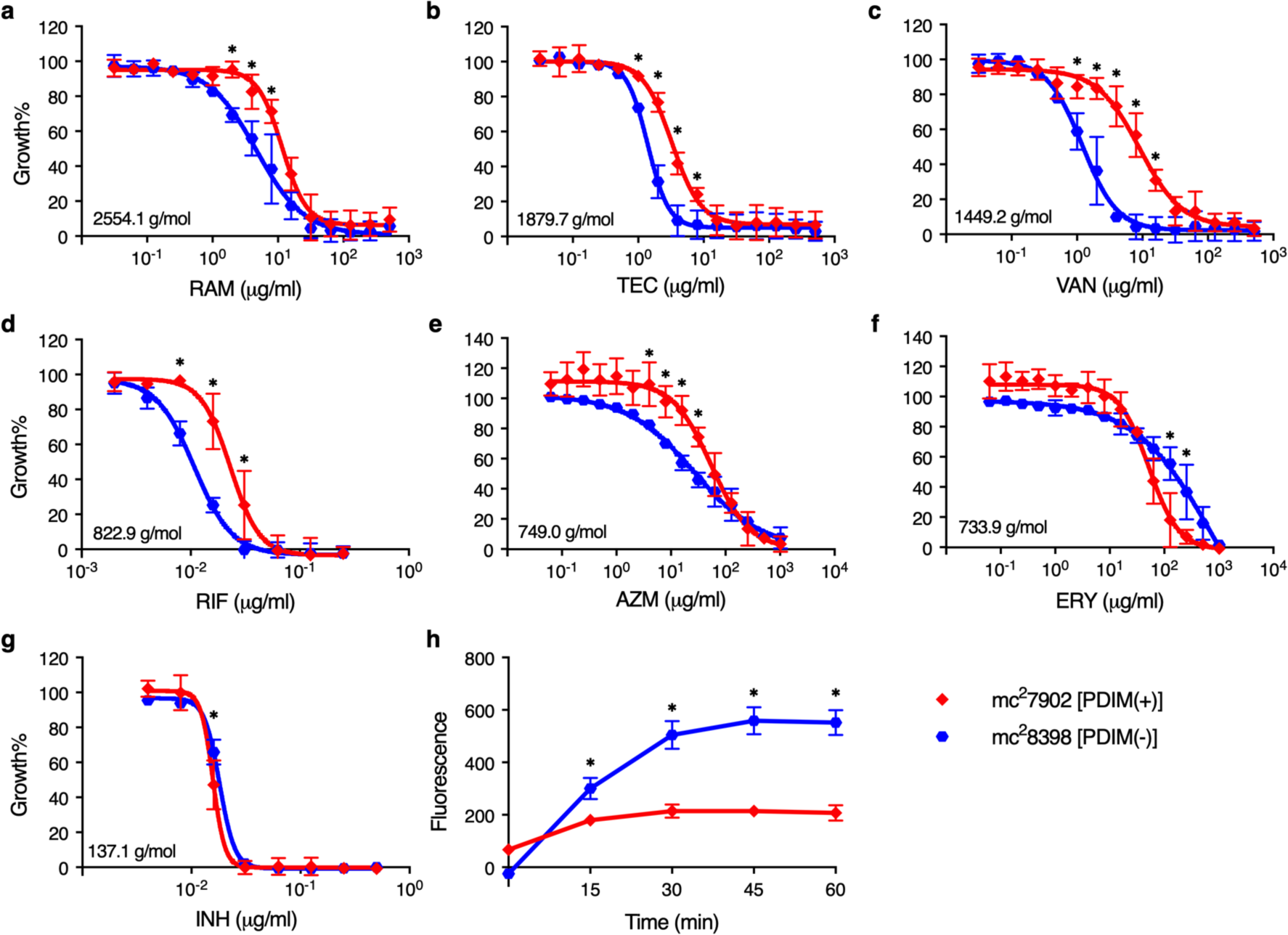
| Resistance of PDIM(-) and PDIM(+) *Mtb* to high molecular weight compounds. **a**–**g**, MIC assays of *Mtb* mc^2^7902 [PDIM(+)] and mc^2^8398 [PDIM(-)] to (**a**) ramoplanin (RAM), (**b**) teicoplanin (TEC), (**c**) vancomycin (VAN), (**d**) rifampicin (RIF), (**e**) azithromycin (AZM), (**f**) erythromycin (ERY), and (**g**) isoniazid (INH). Compounds are arranged by descending molecular weight, which is shown on the MIC plots. MICs were performed in 7H9/OADC/glycerol/tyloxapol + PALM media and bacterial growth was measured after 10 days of incubation and normalized to drug-free controls. Mean ± SD for *n* = 4 biological replicates from two independent experiments. **h**, Ethidium Bromide uptake of mc^2^7902 and mc^2^8398. Uptake in whole cell suspensions was monitored by fluorescence (Ex 355 nm/Em 590 nm). Mean ± SD for *n* = 4 biological replicates, each measured in five technical replicates. Uptake data are representative of two independent experiments. \**P* < 0.001; two-way ANOVA with Šidák’s multiple comparison test.

**Extended Data Fig. 2.**
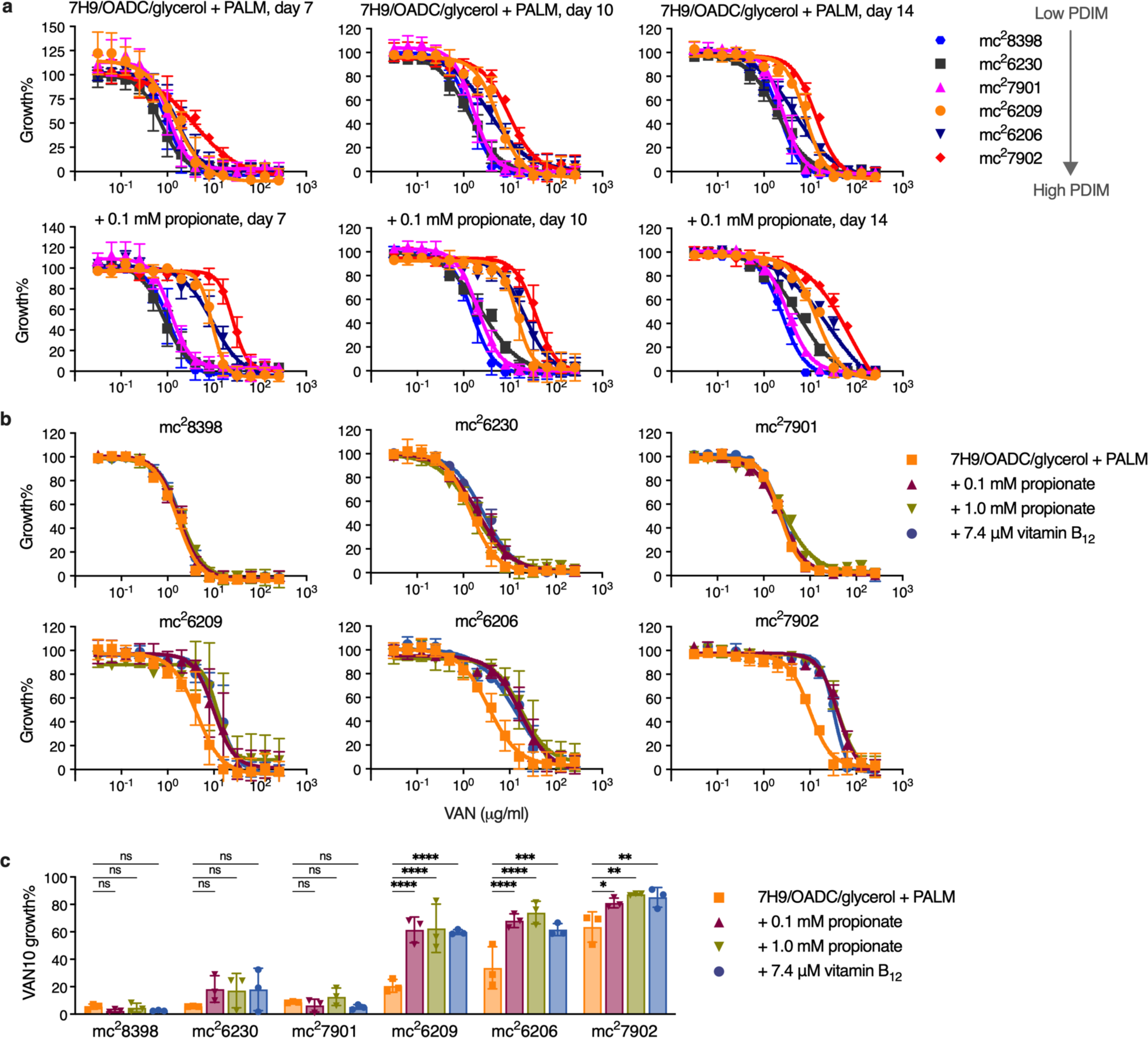
| Propionate and vitamin B_12_ supplementation selectively increase vancomycin resistance of PDIM(+) *Mtb* improving assay robustness and reducing time to result. **a**, Vancomycin MICs for the PDIM reference strain set in standard 7H9/OADC/glycerol/tyloxapol + PALM media and additionally supplemented with 0.1 mM propionate. Growth was measured after 7, 10, and 14 days as indicated. **b**, Vancomycin MICs in standard media and additionally supplemented with 0.1 or 1.0 mM propionate or 7.4 μM vitamin B_12_. Growth was measured after 10 days. Mean ± SD for *n* = 4 biological replicates from two independent experiments. **c**, VAN10 assays in standard and supplemented media. Growth was measured after 10 days. Mean ± SD for *n* = 3 independent experiments, each performed in triplicate. \**P* < 0.05, \*\**P* < 0.01, \*\*\**P* < 0.001, \*\*\*\**P* < 0.0001; two-way ANOVA with Šidák’s multiple comparison test. The day seven data in (**a**) are additionally shown in Fig. 1c and are shown here alongside additional time points. The data in (**b**) includes one of the same experiments shown in (**a**), together with data from an independent experiment. The VAN10-P (+ 0.1 mM propionate) data in (**c**) are additionally shown in Fig. 1e and are shown here alongside additional conditions.

**Extended Data Fig. 3.**
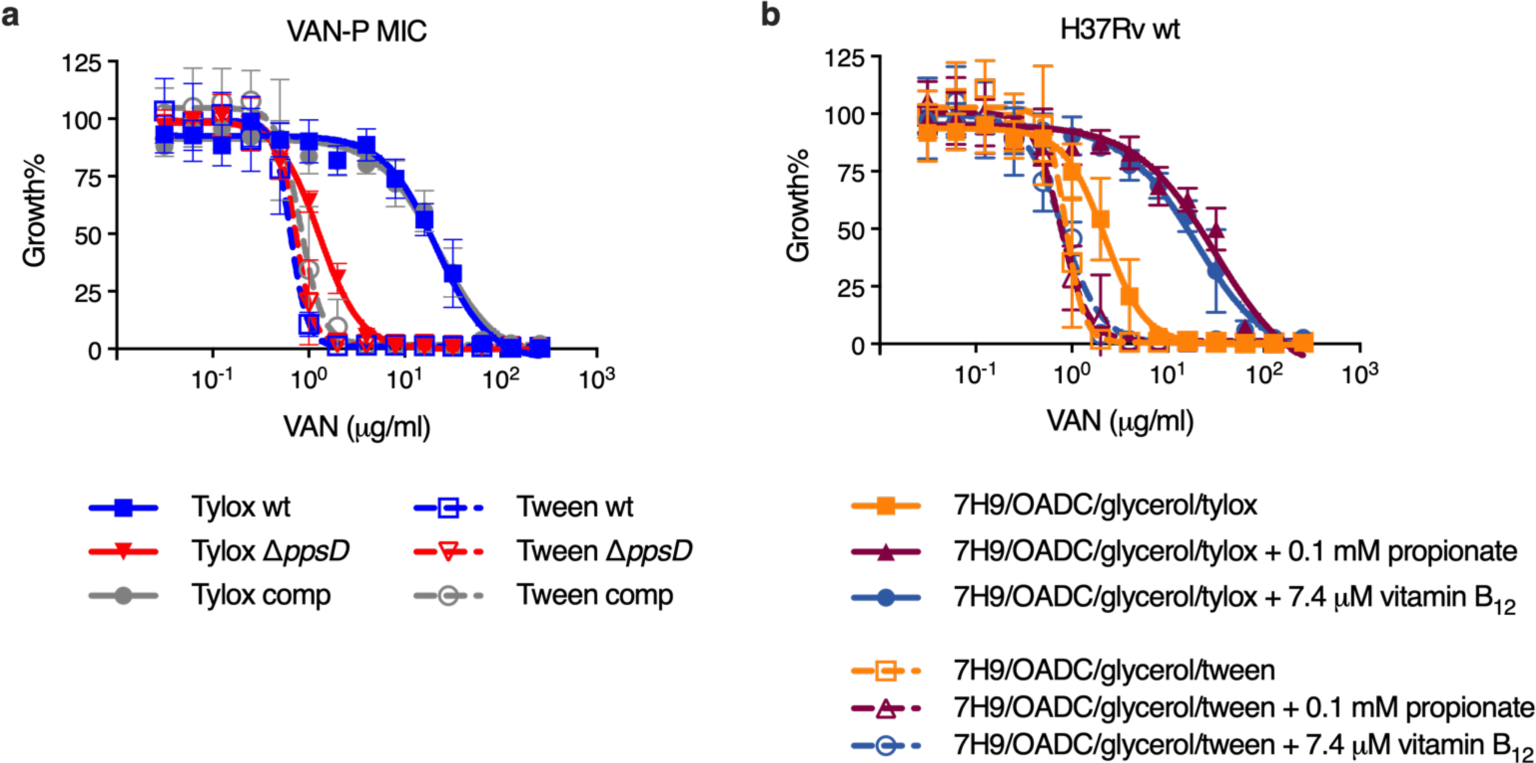
| Tween 80 decreases vancomycin resistance and abolishes PDIM-related differences in MIC. **a**, VAN-P MICs for isogenic PDIM(+) and PDIM(-) *Mtb* H37Rv using either tyloxapol or Tween 80 as the culture detergent. **b**, Vancomycin MICs for PDIM(+) H37Rv wildtype in standard 7H9/OADC/glycerol media and supplemented with propionate or vitamin B_12_ using either tyloxapol or Tween 80 as the detergent. Mean ± SD for *n* = 4 biological replicates from two independent experiments.

**Extended Data Fig. 4.**
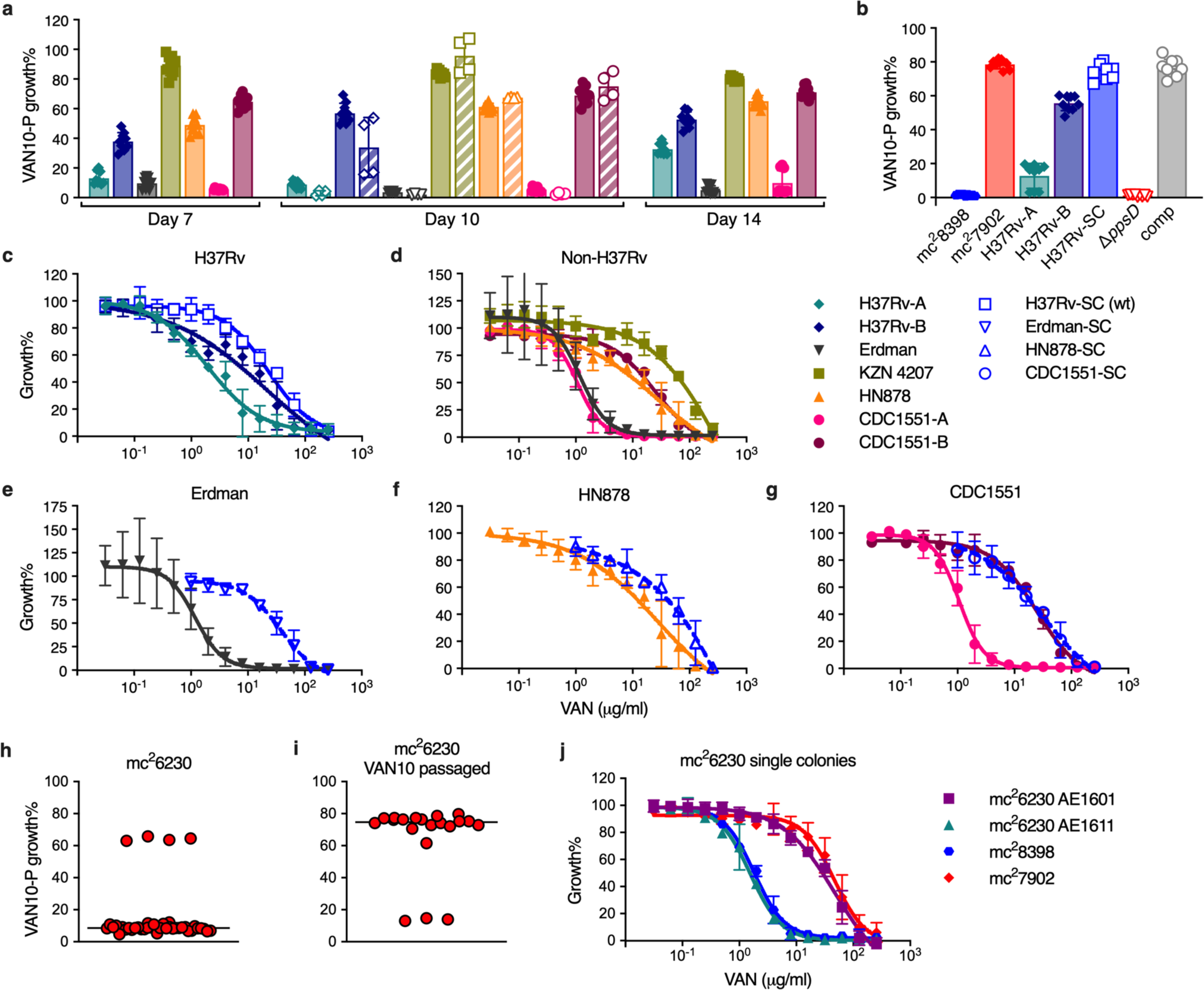
| VAN-P assays predict PDIM levels across different *Mtb* strains and lineages and enable re-isolation of single PDIM(+) clones. **a**, VAN10-P assays for a range of virulent *Mtb* strains belonging to different lineages. Bacterial growth was measured after 7, 10, and 14 days of incubation as indicated. Data are from the same experiment in Fig. 2a and show additional time points plus an independent experimental repeat measured on day 10 (hatched bars with unfilled symbols). **b**, VAN10-P assays for strains with an H37Rv background including mc^2^7902 and mc^2^8398. H37Rv-SC is a single PDIM(+) clone isolated from H37Rv-B by VAN10-P colony screening. This clone was used as our PDIM(+) H37Rv wildtype strain throughout this work and was used to construct H37Rv 11.*ppsD* and 11.*ppsD*::comp isogenic mutants (Supplementary Table 2). Data in (**a**,**b**) show mean ± SD for *n* = 9 pairwise comparisons between triplicate wells, except for the day 10 repeat in (**a**) where *n* = 4 pairwise comparisons between duplicate wells. **c**, VAN-P MICs of H37Rv stocks and H37Rv-SC. **d**, VAN-P MICs of non-H37Rv strains from (**a**). **e**–**g**, VAN-P MICs of single PDIM(+) clones isolated from Rag**^-^**^/**-**^ mice using VAN10-P colony screening for (**e**) Erdman, (**f**) HN878, and (**g**) CDC1551 (see also Extended Data Table 1). Data are plotted together with MIC data from (**d**) for comparison. MIC data in (**c**–**g**) show mean ± SD for *n* = 4 biological replicates from two independent experiments. **h**–**j**, Determination that our *Mtb* mc^2^6230 stock is a mixed population and re-isolation of a single PDIM(+) clone by VAN10-P screening. (**h**) VAN10-P assay of single colonies isolated from our mc^2^6230 stock (*n* = 40) and (**i**) following a single passage in 10 μg/ml vancomycin (*n* = 20). Vancomycin significantly enriched for PDIM(+) bacilli (*P* < 0.0001 two-tailed Mann-Whitney test), facilitating re-isolation of low-frequency PDIM(+) clones. Each colony was assayed in triplicate and data points represent mean VAN10-P growth%. Lines indicate the median. **j**, VAN-P MICs of PDIM(+) (AE1601) and PDIM(-) (AE1611) mc^2^6230 clones identified by VAN10-P colony screening (see also Extended Data Table 1). Mean ± SD for *n* = 6 biological replicates from two independent experiments.

**Extended Data Fig. 5.**
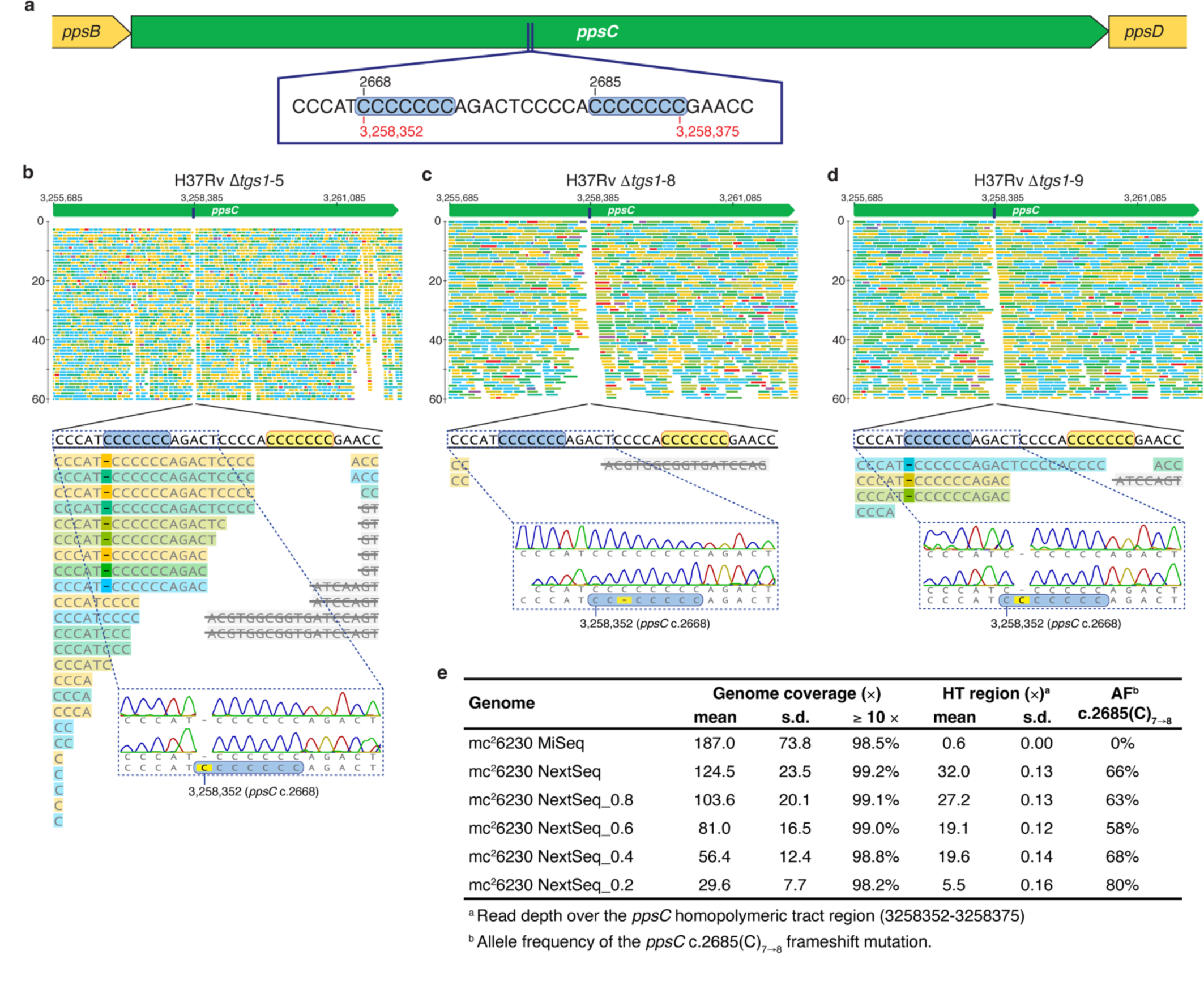
| Assessment of *ppsC* homopolymeric tract mutations. **a**, Schematic showing the location of a homopolymeric tract region in the *ppsC* gene. Sequence inserts show two adjacent 7-cytosine homopolymeric tracts (c.2668 and c.2685) ± 5 bp on either side. Numbers in black indicate the position in the *ppsC* gene and numbers in red the genomic position in the H37Rv genome. **b**–**d**, Analysis of the *ppsC* homopolymeric tract region in 11.*tgs1* mutants and identification of frameshift variants. WGS variant calling failed to identify PDIM mutations in 11.*tgs1*-5, 11.*tgs1*-8 and 11.*tgs1*-9 despite a PDIM(-) result in VAN-P MICs (Fig. 2b) and validation of 11.*tgs1*-9 as PDIM(-) by TLC (Fig. 2c). Close manual inspection of WGS reads showed the *ppsC* homopolymeric tract region is poorly covered by Illumina MiSeq and identified potentially missed variant calls. PCR and Sanger sequencing confirmed the presence of a 2668(C)_7→6_ frameshift mutation in both 11.*tgs1*-5 (**b**) and 11.*tgs1*-9 (**d**) and identified a 2668(C)_7→8_ mutation in 11.*tgs1*-8 that was not covered at all by WGS (**c**). (**b**–**d**) were created with Geneious Prime^®^ 2022.2.2 and Illustrator 26.4.1. Coverage has been cropped to a read depth of 60 ξ. **e**, Identification of an unfixed *ppsC* c.2685(C)_7→8_ frameshift mutation in mc^2^6230 by Illumina NextSeq. VAN-P assays and TLC lipid analysis determined mc^2^6230 is highly PDIM deficient (Fig. 1a,c,e), however, WGS initially failed to identify any PDIM mutations in this strain and we subsequently established our mc^2^6230 stock is a mixed population (Extended Data Fig. 4h). Resequencing using the Illumina NextSeq platform identified an unfixed frameshift mutation in *ppsC* (c.2685(C)_7→8_) that was not detected by Illumina MiSeq due to poor coverage. To assess the relationship between overall coverage and coverage over the homopolymeric region NextSeq reads were randomly downsampled. The number following ‘NextSeq_’ represents the fraction of reads sampled (i.e. 0.8 = 80% of reads retained).

**Extended Data Fig. 6.**
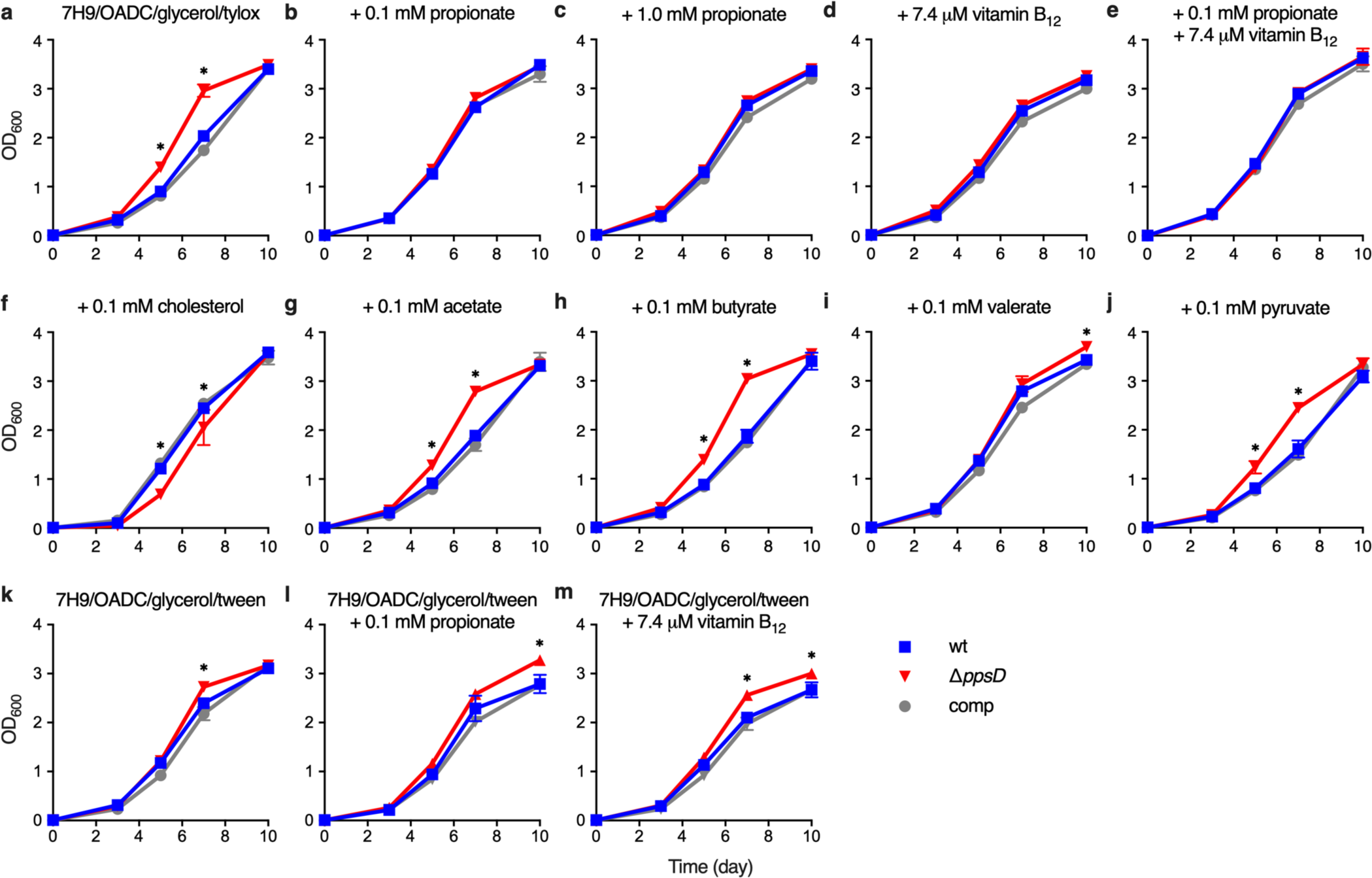
| Effect of different media supplements on growth of PDIM(+) and PDIM(-) *Mtb*. **a**, Growth of PDIM(+) and PDIM(-) *Mtb* H37Rv in standard 7H9/OADC/glycerol/tyloxapol and **b**–**j**, the same media with additional supplements as indicated. **k**–**m**, Growth using Tween 80 instead of tyloxapol as the culture detergent. (**k**) 7H9/OADC/glycerol/Tween 80 and (**l**) the same media supplemented with 0.1 mM propionate, or (**m**) 7.4 μM vitamin B_12_. Mean ± SD for *n* = 3 biological replicates. Data are representative of at least two independent experiments. (**a**,**b**,**d**,**f**) show independent experimental repeats for the conditions in Fig. 3b–e. \**P* < 0.001 for both wt and comp versus 11.*ppsD*; two-way ANOVA with Tukey’s multiple comparison test. For some data points the SD is smaller than the data symbols.

**Extended Data Fig. 7.**
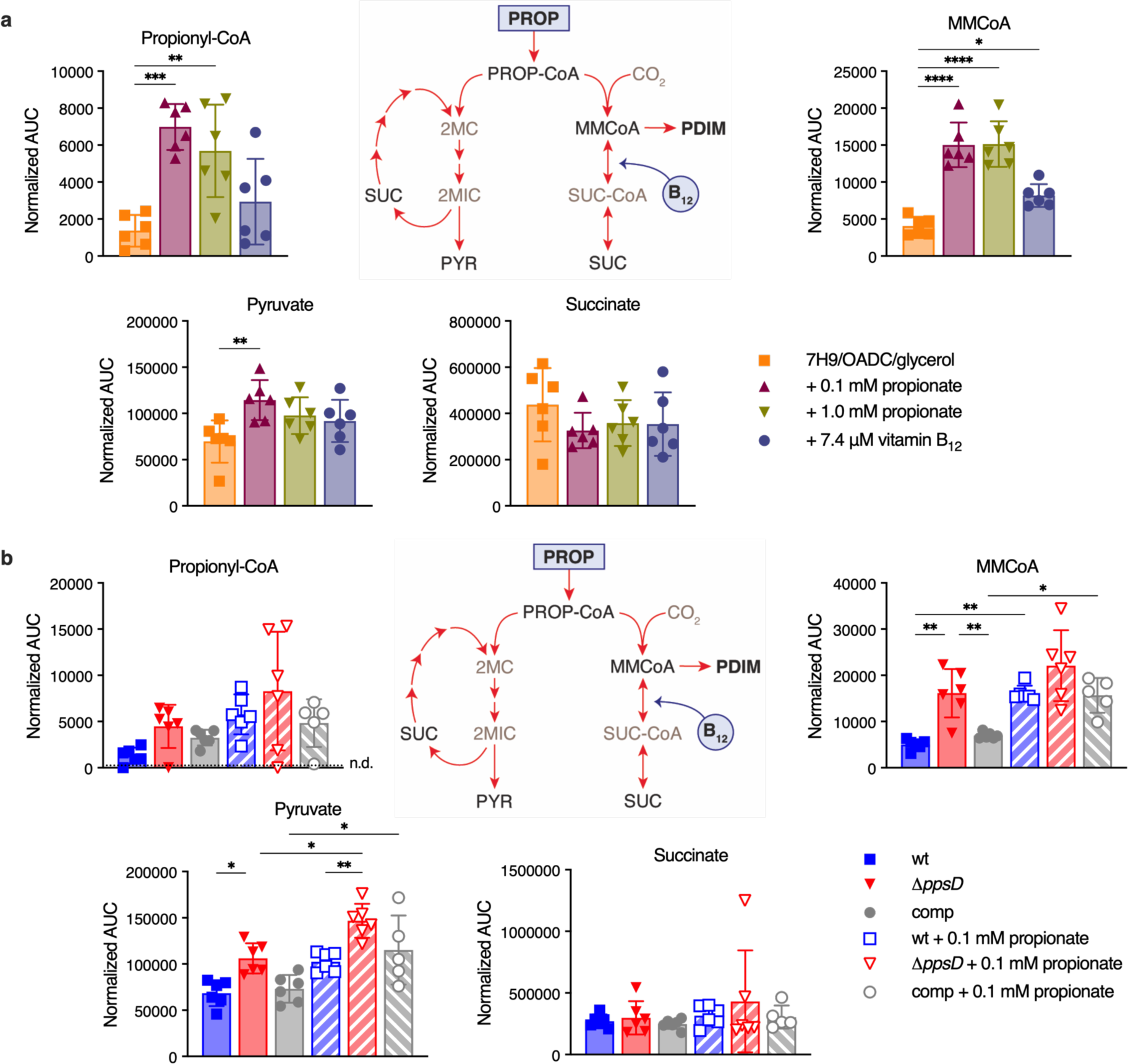
| Effects of propionate and vitamin B_12_ supplementation on MMCoA and propionyl-CoA metabolic pathways in *Mtb*. **a**, Abundance of metabolites in propionyl-CoA and MMCoA metabolism in PDIM(+) *Mtb* H37Rv wildtype grown in standard 7H9/OADC/glycerol/tyloxapol media and supplemented with propionate or vitamin B_12_, and **b**, in PDIM(+) and PDIM(-) H37Rv grown in 7H9/OADC/glycerol/tyloxapol ± 0.1 mM propionate. Abundances are shown as normalized area under the curve (AUC). Mean ± SD for *n* = 6 biological replicates from two independent experiments. \**P* < 0.05, \*\**P* < 0.01, \*\*\**P* < 0.001, \*\*\*\**P* < 0.0001; one-way ANOVA with Tukey’s multiple comparison test. Significant differences compared to unsupplemented media are indicated in (**a**), and between ± propionate for each strain and between strains for each condition in (**b**). PROP, propionate; PROP-CoA, propionyl-CoA; MMCoA, methylmalonyl-CoA; SUC-CoA, succinyl-CoA; SUC, succinate; 2MC/2MIC, 2-methyl(iso)citrate; and PYR, pyruvate. Succinyl-CoA and methyl(iso)citrate were not able to be detected by our method. Propionyl-CoA was close to the detection limit and was not detected in all samples (n.d. = not detected). The data for MMCoA are also shown in Fig. 3f,g.

**Extended Data Fig. 8.**
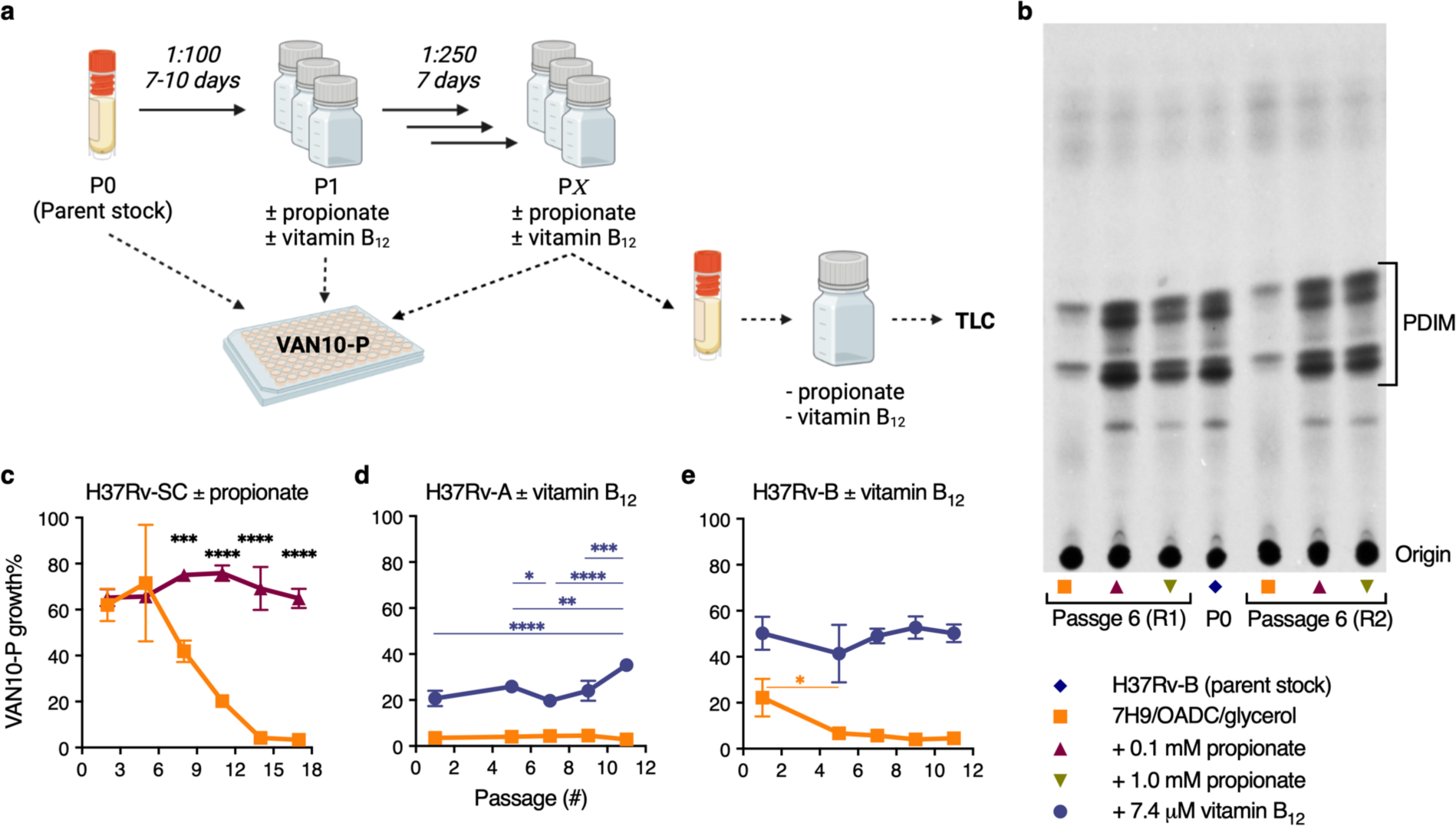
| Propionate and vitamin B_12_ supplementation prevent PDIM loss in *Mtb*. **a**, Schematic overview of *in vitro* evolution experiments. Triplicate inkwells containing standard 7H9/OADC/glycerol/tyloxapol or media supplemented with propionate or vitamin B_12_ were inoculated with frozen *Mtb* culture stock (P0) and incubated for 7-10 days (P1). Cultures were then diluted into fresh media every 7 days for serial passage (P2 to P*X*). Selected passages were input into VAN10-P assays at the time of passage to assess PDIM production over the course of the experiment. For TLC lipid analysis, frozen stocks were first outgrown in media without propionate or vitamin B_12_ for a single passage to allow the strains to recover before ^14^C-labelling. Figure created with BioRender.com. **b**, TLC lipid analysis of H37Rv-B before and after six serial passages in ± 0.1 or 1.0 mM propionate. This figure shows the full TLC plate from Fig. 4a with results for both biological replicates analysed by TLC (R2 is shown in Fig. 4a). **c**, VAN10-P assays for H37Rv-SC [PDIM(+) H37Rv wildtype] passaged in ± 0.1 mM propionate. **d**, H37Rv-A and **e**, H37Rv-B passaged in ± 7.4 μM vitamin B_12_. Mean ± SD for *n* = 3 biological replicates, each assayed in triplicate. \**P* < 0.05, \*\**P* < 0.01, \*\*\**P* < 0.001, \*\*\*\**P* < 0.0001; two-way ANOVA with Šidák’s (**c**) or Tukey’s (**d**,**e**) multiple comparison test. Significant differences between conditions are indicated in (**c**) and between timepoints in (**d**,**e**).

