## Supplemental tables and figures for "The PDIM paradox of *Mycobacterium tuberculosis*: new solutions to a persistent problem"

**This PDF file includes:**

**Supplementary Figures 1–6**

Supplementary Fig. 1. Vancomycin resistance is predictive of PDIM production in *Mtb*.
Supplementary Fig. 2. VAN10 assay development.
Supplementary Fig. 3. VAN-P MIC assay validation in *Mtb* CDC1551.
Supplementary Fig. 4. Effect of leucine supplementation on vancomycin sensitivity and growth of PDIM(+) and PDIM(-) *Mtb*
Supplementary Fig. 5. Effect of propionate supplementation on the sensitivity of PDIM(+) and PDIM(-) *Mtb* to compounds with different modes of action and molecular weights
Supplementary Fig. 6. Propionate increases the resistance of *Mtb* CDC1551 to rifampicin, but not isoniazid, in a PDIM-dependent manner.

**Supplementary Tables 1–8**

Supplementary Table 1. BSL2-approved attenuated *Mtb* strains included in this study.
Supplementary Table 2. Virulent *Mtb* strains included in this study.
Supplementary Table 3. Culture media reagents and supplements used in this study.
Supplementary Table 4. Primers used in this study.
Supplementary Table 5. Inhibitors used in this study.
Supplementary Table 6. Chemical standards used for LC-MS.
Supplementary Table 7. Putative metabolites used for LC-MS data normalization.
Supplementary Table 8. Whole genome sequencing accession numbers.

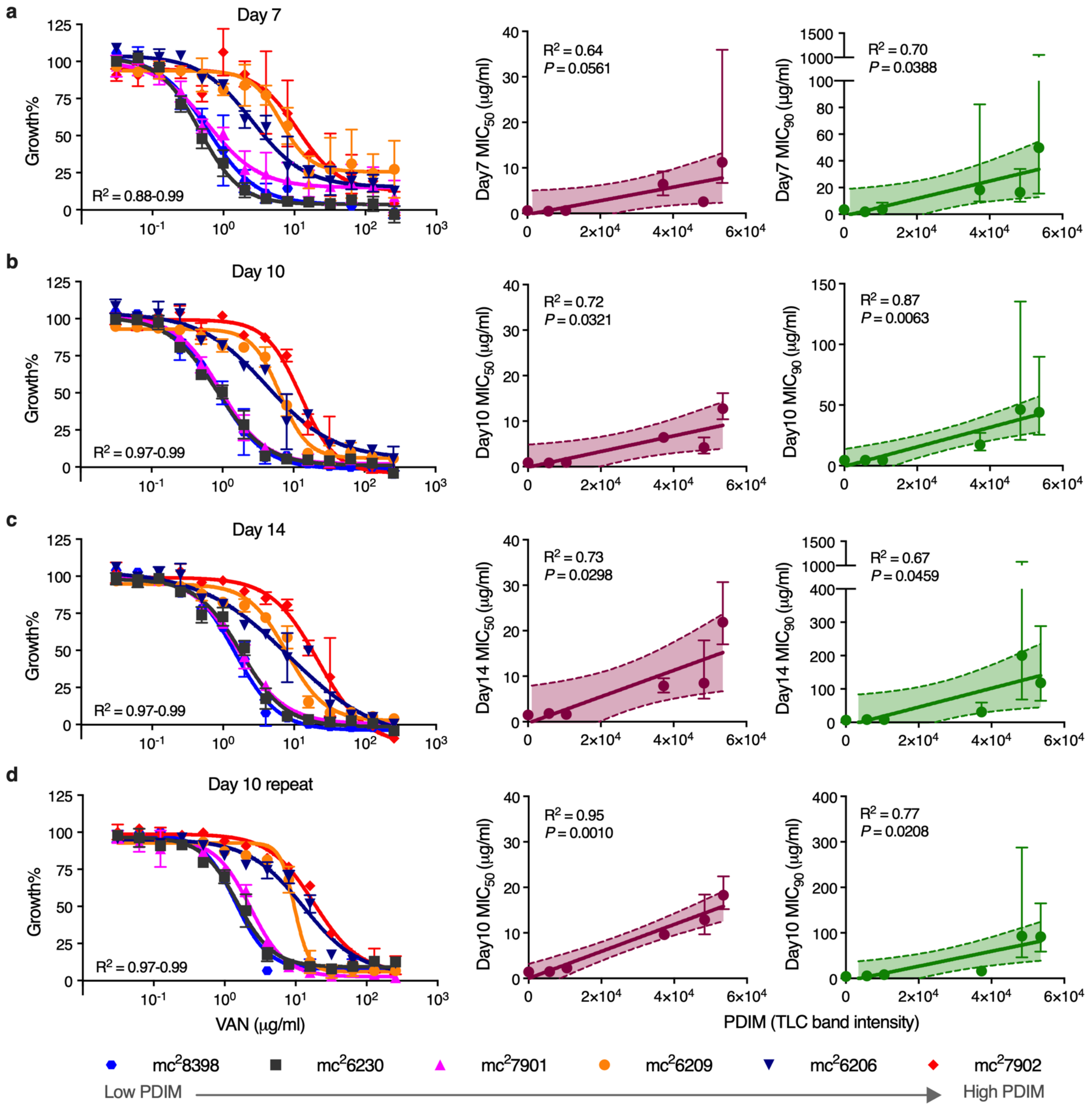


**Supplementary Fig. 1 | Vancomycin resistance is predictive of PDIM production in *Mtb*. a–c**, Vancomycin resistance of six BSL2-approved attenuated *Mtb* H37Rv strains (Supplementary Table 1) and correlation with PDIM lipid levels determined by TLC (Fig. 1a). These strains comprise our PDIM reference strain set. Minimum inhibitory concentration (MIC) assays were performed using the broth microdilution method in 7H9/OADC/glycerol/tyloxapol media with pantothenate, arginine, leucine, and methionine (PALM) supplements. Bacterial growth was measured after (**a**) 7, (**b**) 10, and (**c**) 14 days of incubation, and normalized to drug-free control wells. Simple linear regression was performed to assess the correlation between vancomycin $MIC_{50}$ (maroon) and $MIC_{90}$ (green) with PDIM levels as determined by TLC densitometric analysis of Fig. 1a. **d**, Experimental repeat measured after 10 days of incubation. MIC plots show mean ± SD for $n = 2$ technical replicates from a single experiment. Bands on regression plots show the 95% CI and error bars show the 95% CI for MIC values calculated from the curve fit.

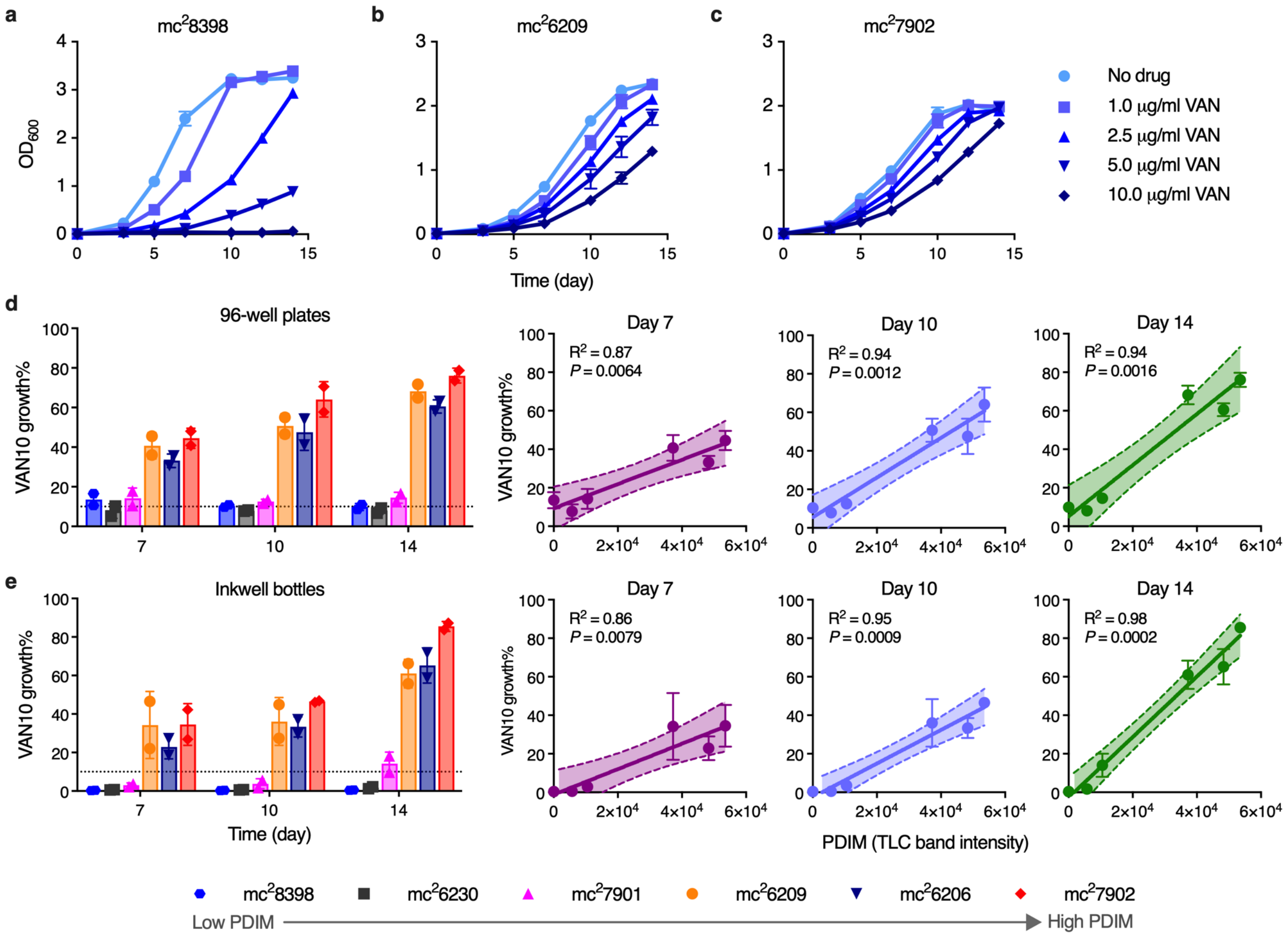


**Supplementary Fig. 2 | VAN10 assay development. a–c**, Growth curves of (**a**) *Mtb* mc$^2$8398 [PDIM(-)], (**b**) mc$^2$6209 [PDIM(+)], and (**c**) mc$^2$7902 [PDIM(+)] (see Fig. 1a) in standard 7H9/OADC/glycerol/tyloxapol + PALM media with increasing concentrations of vancomycin. Mean ± SD for $n = 3$ biological replicates. For some data points the SD is smaller than the data symbols. This experiment was performed once. **d**,**e**, 'VAN10 assay' showing relative growth of the PDIM reference strain set in standard 7H9/OADC/glycerol/tyloxapol + PALM media with 10 μg/ml vancomycin compared to drug-free controls in (**d**) 96-well plates and (**e**) inkwell bottles. Bacterial growth was assessed by measuring optical density after 7, 10, and 14 days of incubation as indicated. VAN10 OD / VAN0 OD × 100 = VAN10 growth%. Simple linear regression was performed to assess the correlation between VAN10 growth% at each time point and PDIM lipid levels as determined by TLC densitometric analysis of Fig. 1a. Mean ± SD for $n = 2$ independent experiments, each performed in triplicate. Bands on regression plots show the 95% CI.

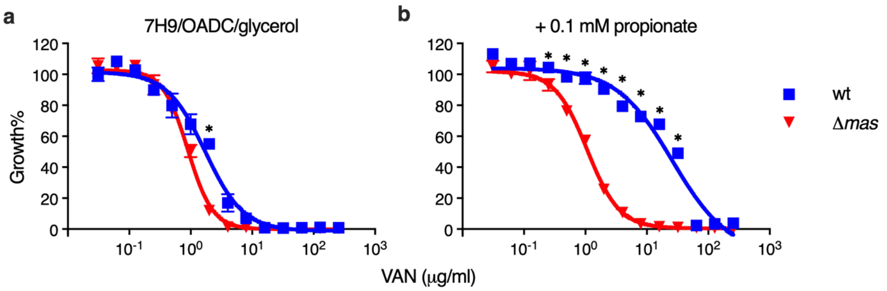


**Supplementary Fig. 3 | VAN-P MIC assay validation in *Mtb* CDC1551.** Vancomycin MIC assays of isogenic PDIM(+) (wildtype) and PDIM(-) (Δ*mas*) CDC1551 in **a**, standard 7H9/OADC/glycerol/tyloxapol media and **b**, supplemented with 0.1 mM propionate. Mean ± SD for $n = 2$ technical replicates from a single experiment. This experiment was performed once. $P < 0.001$; two-way ANOVA with Šidák's multiple comparison test. For some data points the SD is smaller than the data symbols.

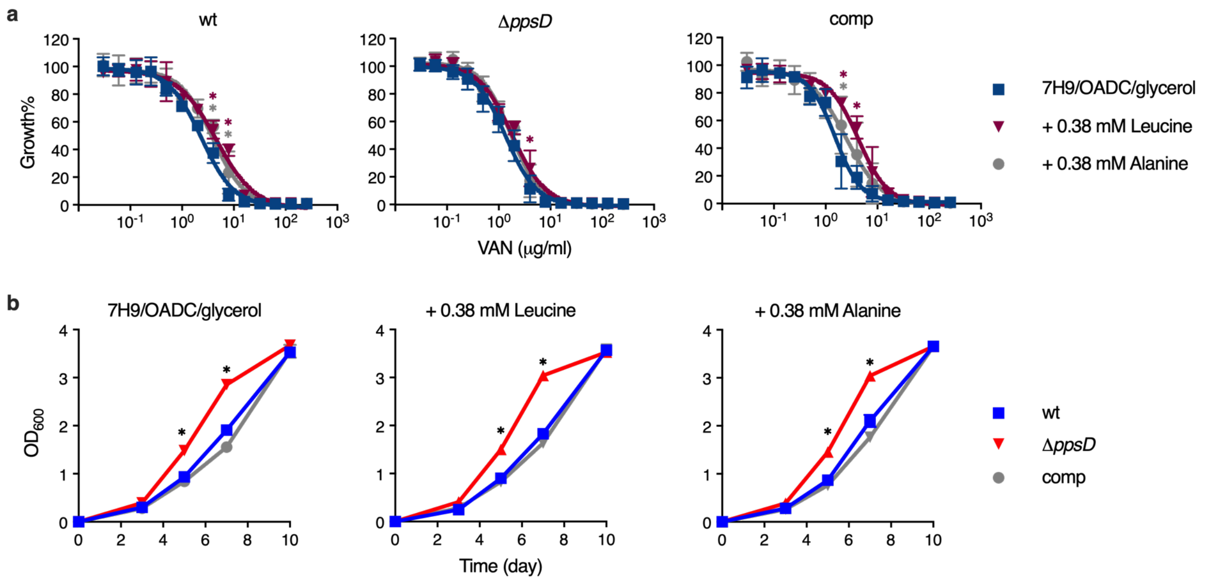


**Supplementary Fig. 4 | Effect of leucine supplementation on vancomycin sensitivity and growth of PDIM(+) and PDIM(-) *Mtb*. a**, Vancomycin MIC assays of isogenic PDIM(+) and PDIM(-) *Mtb* H37Rv in standard 7H9/OADC/glycerol/tyloxapol media and supplemented with either 0.38 mM L-leucine (50 mg/l, same concentration provided in PALM supplemented media) or 0.38 mM L-alanine as a non-branched-chain amino acid control. Mean ± SD for $n$ = 4 biological replicates from two independent experiments. $P < 0.001$ compared to unsupplemented media; two-way ANOVA with Tukey's multiple comparison test. **b**, Growth of PDIM(+) and PDIM(-) H37Rv in standard 7H9/OADC/glycerol/tyloxapol media and supplemented with either L-leucine or L-alanine. Mean ± SD for $n$ = 3 biological replicates. *$P < 0.001$ for both wt and comp versus Δ*ppsD*; two-way ANOVA with Tukey's multiple comparison test. This experiment was repeated once with similar results. For some data points the SD is smaller than the data symbols.

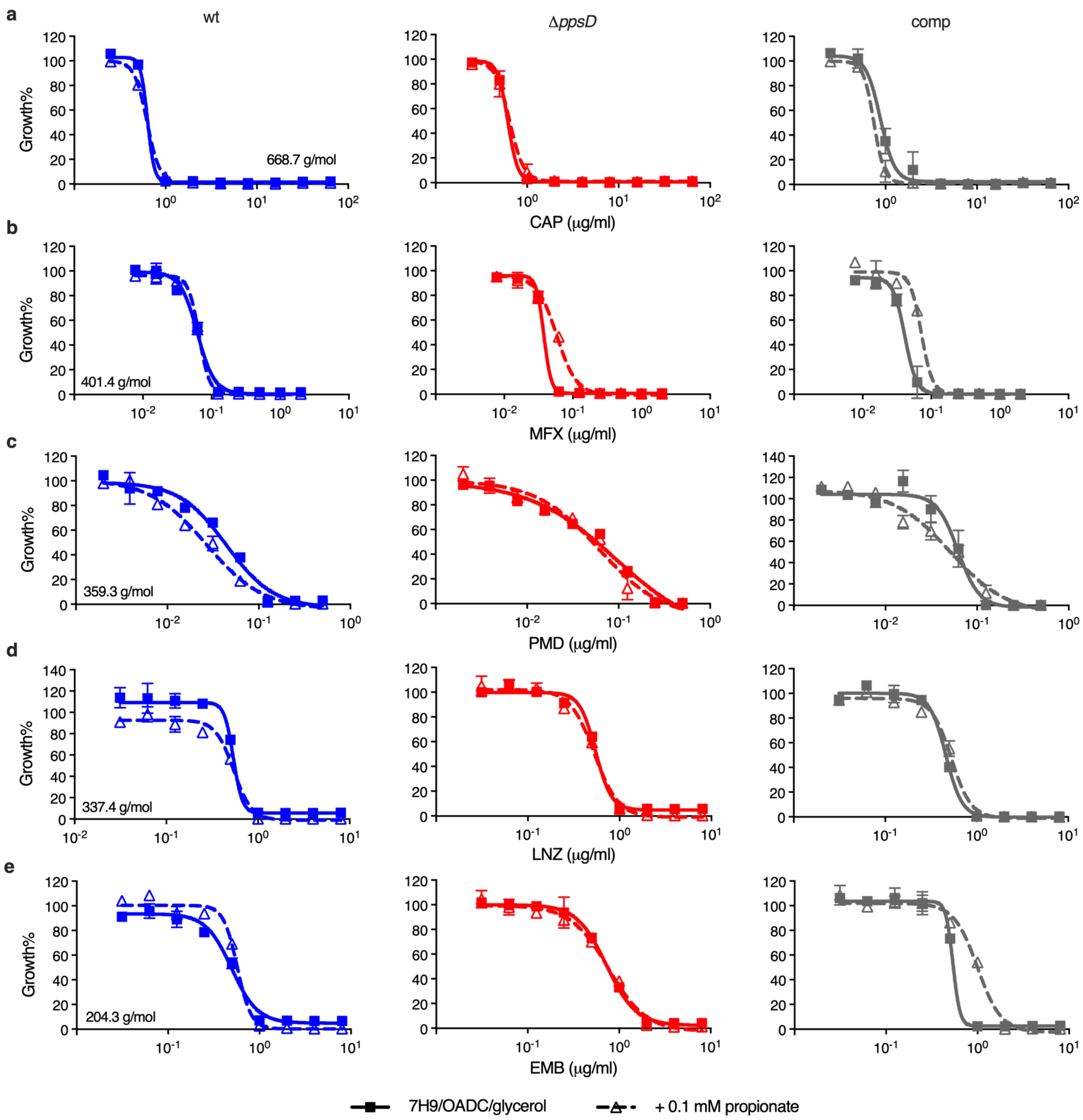


**Supplementary Fig. 5 | Effect of propionate supplementation on the sensitivity of PDIM(+) and PDIM(-) *Mtb* to compounds with different modes of action and molecular weights.** MIC assays of isogenic PDIM(+) and PDIM(-) *Mtb* H37Rv in 7H9/OADC/glycerol/tyloxapol media ± 0.1 mM propionate for **a**, capreomycin (CAP); **b**, moxifloxacin (MFX); **c**, pretomanid (PMD); **d**, linezolid (LNZ); and **e**, ethambutol (EMB). The molecular weight of each compound is shown on the corresponding MIC plot. Mean ± SD for $n = 2$ technical duplicates from a single experiment. This experiment was performed once. For some data points the SD is smaller than the data symbols.

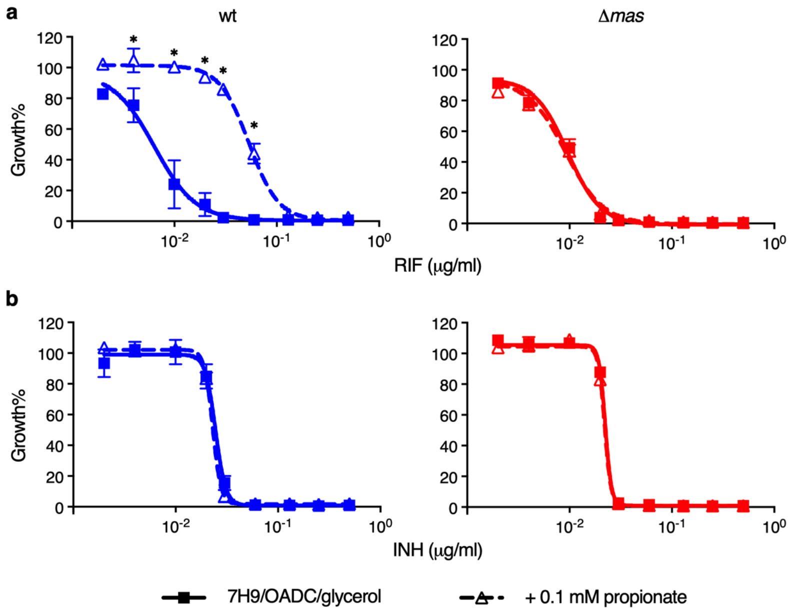


**Supplementary Fig. 6 | Propionate increases the resistance of *Mtb* CDC1551 to rifampicin, but not isoniazid, in a PDIM-dependent manner.** Sensitivity of isogenic PDIM(+) and PDIM(-) CDC1551 in 7H9/OADC/glycerol/tyloxapol media ± 0.1 mM propionate to **a**, rifampicin (RIF), and **b**, isoniazid (INH). Mean ± SD for $n = 2$ technical replicates from a single experiment. This experiment was performed once. $^*P < 0.001$; two-way ANOVA with Šidák's multiple comparison test. For some data points the SD is smaller than the data symbols.

**Supplementary Table 1 | BSL2-approved attenuated *Mtb* strains included in this study.** These strains comprise our PDIM reference strain set, with the exception of mc$^2$6230 AE1601 and AE1611, which are PDIM(+) and PDIM(-) clones isolated from our mc$^2$6230 stock.

| Strain name | Genotype | Reference |
|---|---|---|
| mc$^2$6206 | H37Rv Δ*panCD* Δ*leuCD* | Jain *et al.* (2014)[28] |
| mc$^2$6209 | H37Rv Δ*panCD* Δ*leuCD* Δ*secA2* | Jain *et al.* (2014)[28] |
| mc$^2$6230 | H37Rv ΔRD1 Δ*panCD* | Sambandamurthy *et al.* (2006)[65]; Jain *et al.* (2012)[66] |
| mc$^2$7901 | H37Rv Δ*panCD* Δ*leuCD* Δ*metA* | Vilchèze *et al.* (2018)[67] |
| mc$^2$7902 | H37Rv Δ*panCD* Δ*leuCD* Δ*argB* | Vilchèze *et al.* (2018)[67] |
| mc$^2$8398 | H37Rv Δ*metA* Δ*argB* | This study |
| mc$^2$6230 AE1601* | H37Rv ΔRD1 Δ*panCD* [PDIM(+)] | This study |
| mc$^2$6230 AE1611† | H37Rv ΔRD1 Δ*panCD* [PDIM(-)] | This study |

*PDIM(+) and †PDIM(-) clones identified by VAN10-P screening of single colonies (see Extended Data Fig. 4).

**Supplementary Table 2 | Virulent *Mtb* strains included in this study.** Single PDIM(+) clones isolated in this study have been assigned unique 'AE' reference numbers as indicated.

| Strain name | Genotype | Source/Construction |
|---|---|---|
| H37Rv-A | H37Rv | In-house culture collection |
| H37Rv-B | H37Rv | C57BL/6 mouse passage of H37Rv-A |
| H37Rv-SC (AE1028)* | H37Rv | PDIM(+) clone isolated from H37Rv-B |
| H37Rv Δ*ppsD* | H37Rv Δ*Rv2934* hyg$^R$ | Specialized transduction of H37Rv-SC |
| H37Rv Δ*ppsD*::comp | H37Rv Δ*Rv2934*::comp hyg$^R$ kan$^R$ | Transformation of H37Rv Δ*ppsD* with pMV361-*ppsD* |
| H37Rv Δ*tgs1*-1 | H37Rv Δ*Rv3130c* hyg$^R$ | Specialized transduction of H37Rv-B |
| H37Rv Δ*tgs1*-2 | H37Rv Δ*Rv3130c* hyg$^R$ | Specialized transduction of H37Rv-B |
| H37Rv Δ*tgs1*-3 | H37Rv Δ*Rv3130c* hyg$^R$ | Specialized transduction of H37Rv-B |
| H37Rv Δ*tgs1*-4 | H37Rv Δ*Rv3130c* hyg$^R$ | Specialized transduction of H37Rv-B |
| H37Rv Δ*tgs1*-5 | H37Rv Δ*Rv3130c* hyg$^R$ | Specialized transduction of H37Rv-B |
| H37Rv Δ*tgs1*-7 | H37Rv Δ*Rv3130c* hyg$^R$ | Specialized transduction of H37Rv-B |
| H37Rv Δ*tgs1*-8 | H37Rv Δ*Rv3130c* hyg$^R$ | Specialized transduction of H37Rv-B |
| H37Rv Δ*tgs1*-9 | H37Rv Δ*Rv3130c* hyg$^R$ | Specialized transduction of H37Rv-B |
| CDC1551-A | CDC1551 | In-house culture collection |
| CDC1551-B | CDC1551 | In-house culture collection |
| CDC1551 Δ*mas* | CDC1551 Δ*Rv2940c* hyg$^R$ | Specialized transduction of CDC1551-B |
| Erdman | Erdman | In-house culture collection |
| HN878 | HN878 | In-house culture collection |
| KZN 4207 | KZN 4207 | In-house culture collection |
| CDC1551-SC (AE5005)* | CDC1551 | PDIM(+) clone isolated from a Rag-/- mouse |
| Erdman-SC (AE3003)* | Erdman | PDIM(+) clone isolated from a Rag-/- mouse |
| HN878-SC (AE8005)* | HN878 | PDIM(+) clone isolated from a Rag-/- mouse |

*PDIM(+) clone identified by VAN10-P screening of single colonies (see Extended Data Fig. 4).

### Supplementary Table 3 | Culture media reagents and supplements used in this study.

| Name | Catalog # | Supplier |
|---|---|---|
| Middlebrook 7H9 | 271310 | BD Difco |
| Middlebrook 7H10 | 262710 | BD Difco |
| Glycerol | G33 | Fisher Chemical |
| Dextrose (D-glucose) | D16 | Fisher Chemical |
| Tyloxapol | T8761 | Sigma-Aldrich |
| Tween 80 | P1754 | Sigma-Aldrich |
| Sodium oleate | 11-1280 | Strem Chemicals |
| Bovine serum albumin fraction V | A-420-1 | GoldBio |
| Catalase | C1345 | Sigma-Aldrich |
| D-Calcium pantothenate | 243305000 | Acros Organics |
| L-Arginine | A5006 | Sigma-Aldrich |
| L-Leucine | L8000 | Sigma-Aldrich |
| L-Methionine | 166165000 | Acros Organics |
| Hygromycin B | 10687010 | Invitrogen |
| Kanamycin | BP906 | Fisher BioReagents |
| Sodium propionate | P1880 | Sigma-Aldrich |
| Vitamin $B_{12}$ | V2876 | Sigma-Aldrich |
| Sodium pyruvate | BP356 | Fisher BioReagents |
| Sodium acetate | A13184.30 | Thermo Scientific Chemicals |
| Sodium butyrate | A11079.06 | Thermo Scientific Chemicals |
| Valeric acid | 149570050 | Acros Organics |
| Cholesterol | C3045 | Sigma-Aldrich |

### Supplementary Table 4 | Primers used in this study.

| Primer name | Sequence |
|---|---|
| **ppsC homopolymeric tract region** | |
| ppsC_homopoly_F | CTGCTGACCCACTCGATCA |
| ppsC_homopoly_R | TGTCGGGCCTGTGGTAAG |
| **Knockout strain confirmation** | |
| Universal_uptag | GATGTCTCACTGAGGTCTCT |
| Rv2940c_LL | TCATGGGCATCACGTTAC |
| Rv2940c_LR | AGAACTCAGCATCGAAACC |
| Rv2934_LL | CATGCAGGCGGTACTCAC |
| Rv2934_LR | GGATCGGGATCGTAGAAC |
| Rv3130c_LL | CACAATGGCTTTCATTCG |
| Rv3130c_LL | GGGTACAGGGACGTAGGC |
| **pMV361-*ppsD* vector construction and confirmation** | |
| pMV361_ppsD_Hsp60_for | GATCCAGCTGCAGAATTCATGACAAGTCTGGCGGAGCG |
| pMV361_ppsD_rev | CTACGTCGACATCGATAAGCTTCTTATCCTCCCTGTACTTCAGGTTTAGGT |
| pMV361_vector_for | GAATTCTGCAGCTGGATCCGC |
| complement_vector_rev | GAAGCTTATCGATGTCGACGTAG |
| pMV361-LL | GGAATCACTTCGCAATGGC |
| pMV361-LR | ATCGTACGCTAGTTAACTACG |
| ppsD_sequencing_2 | CGCCAGCGCTCGCAAC |
| ppsD_sequencing_3 | CGGTGCTGGCCGACTGGATG |
| ppsD_sequencing_4 | GGCATAAAAATCGGCCGGCG |
| ppsD_sequencing_5 | TAATTCGCGGCGGGGACAGC |
| ppsD_sequencing_6 | CGAAGCGGCGGAAGAGAATCC |
| ppsD_sequencing_7 | TCTGGGGGCCAATTGCGCGG |

### Supplementary Table 5 | Inhibitors used in this study.

| Name | Abbreviation | Catalog # | Supplier |
|---|---|---|---|
| Azithromycin dihydrate | AZM | A2076 | TCI America |
| Bedaquiline | BDQ | Gift | Kevin Pethe, Nanyang Technological University, Singapore |
| Capreomycin sulfate | CAP | J66684 | Alfa Aesar |
| Ethambutol dihydrochloride | EMB | J60695 | Alfa Aesar |
| Erythromycin | ERY | E0751 | TCI America |
| Isoniazid | INH | I3377 | Sigma |
| Linezolid | LNZ | 460592500 | Acros Organics |
| Moxifloxacin hydrochloride | MFX | 457960010 | Acros Organics |
| Pretomanid | PMD | HY-10844 | MedChemExpress |
| Ramoplanin | RAM | sc-253424 | Santa Cruz Biotechnology |
| Rifampicin | RIF | R3501 | Sigma |
| Teicoplanin (A2) | TEC | RS048 | BIOTANG Inc. |
| Vancomycin hydrochloride | VAN | J62790 | Alfa Aesar |

### Supplementary Table 6 | Chemical standards used for LC-MS.

| Name | Catalog # | Supplier |
|---|---|---|
| n-Propionyl coenzyme A lithium salt | P5397 | Sigma-Aldrich |
| Methylmalonyl coenzyme A tetralithium salt | sc-215385 | ChemCruz |
| Succinic acid | 33272 | Alfa Aesar |
| Sodium pyruvate | BP356 | Fisher BioReagents |

### Supplementary Table 7 | Putative metabolites used for LC-MS data normalization.

| Compound | m/z | Pathway |
|---|---|---|
| Glycine | 74.0248 | Amino acids and amino acid metabolism |
| Pyruvic acid | 87.0088 | Central carbon |
| L-Alanine | 88.0404 | Amino acids and amino acid metabolism |
| L-Serine | 104.0353 | Amino acids and amino acid metabolism |
| Uracil | 111.0200 | Nucleotide metabolism |
| L-Proline | 114.0561 | Amino acids and amino acid metabolism |
| L-Valine | 116.0717 | Amino acids and amino acid metabolism |
| Succinic acid | 117.0193 | Central carbon |
| L-Threonine | 118.0510 | Amino acids and amino acid metabolism |
| Nicotinic acid | 122.0248 | Nicotinate and nicotinamide metabolism |
| L-Leucine | 130.0874 | Amino acids and amino acid metabolism |
| L-Isoleucine | 130.0874 | Amino acids and amino acid metabolism |
| L-Aspartic Acid | 132.0302 | Amino acids and amino acid metabolism |
| L-Malic acid | 133.0142 | Central carbon |
| Adenine | 134.0472 | Nucleotide metabolism |
| Salicylic acid | 137.0244 | Other |
| 4-Hydroxybenzoic acid | 137.0244 | Folate biosynthesis |
| alpha-Ketoglutaric acid | 145.0142 | Central carbon |
| L-Glutamine | 145.0619 | Amino acids and amino acid metabolism |
| L-Lysine | 145.0983 | Amino acids and amino acid metabolism |
| L-Glutamic acid | 146.0459 | Amino acids and amino acid metabolism |
| L-Methionine | 148.0438 | Amino acids and amino acid metabolism |
| Guanine | 150.0421 | Nucleotide metabolism |
| L-Histidine | 154.0622 | Amino acids and amino acid metabolism |
| Orotic acid | 155.0098 | Nucleotide metabolism |
| L-Phenylalanine | 164.0717 | Amino acids and amino acid metabolism |
| DL-Glyceraldehyde 3-phosphate | 168.9907 | Central carbon |
| Dihydroxyacetone phosphate | 168.9907 | Central carbon |
| D-Glycerol 3-phosphate | 171.0064 | Central carbon |
| Shikimic acid | 173.0455 | Amino acids and amino acid metabolism |
| N2-Acetyl-L-ornithine | 173.0932 | Amino acids and amino acid metabolism |
| L-Arginine | 173.1044 | Amino acids and amino acid metabolism |
| a-D-Glucose | 179.0561 | Central carbon |
| L-Tyrosine | 180.0666 | Amino acids and amino acid metabolism |
| L-Tryptophan | 203.0826 | Amino acids and amino acid metabolism |
| Thymidine | 241.0830 | Nucleotide metabolism |
| D-Fructose 6-P/a-D-Glucose 6-P | 259.0224 | Central carbon |
| Adenosine | 266.0895 | Nucleotide metabolism |
| Inosine | 267.0735 | Nucleotide metabolism |
| dTMP | 321.0493 | Nucleotide metabolism |
| cAMP | 328.0452 | Nucleotide metabolism |
| Trehalose/Maltose | 341.1089 | Central carbon |
| 5'-AMP | 346.0558 | Nucleotide metabolism |
| dTDP | 401.0157 | Nucleotide metabolism |
| Trehalose 6-phosphate | 421.0753 | Central carbon |
| ADP | 426.0221 | Nucleotide metabolism |
| ATP | 505.9885 | Nucleotide metabolism |
| Acetyl-CoA | 808.1185 | Central carbon |
| Propionyl-CoA | 822.1341 | Central carbon |
| Methylmalonyl-CoA | 866.1240 | Central carbon |

**Supplementary Table 8 | Whole genome sequencing accession numbers.** BioProject PRJNA923717.

| Strain | BioSample accession | SRA accession number/s | | |
|---|---|---|---|---|
| mc$^2$6206 | SAMN32734011 | SRR23080350 | | |
| mc$^2$6209 | SAMN32734012 | SRR23080349 | | |
| mc$^2$6230 | SAMN32734013 | SRR23080327 | SRR23080338* | SRR23080318† |
| mc$^2$6230 AE1601 | SAMN35027541 | SRR24495990† | | |
| mc$^2$6230 AE1611 | SAMN35027542 | SRR24495989 | | |
| mc$^2$7901 | SAMN32734014 | SRR23080317 | | |
| mc$^2$7902 | SAMN32734015 | SRR23080316 | | |
| mc$^2$8398 | SAMN32734016 | SRR23080315 | | |
| H37Rv-A | SAMN32734017 | SRR23080314 | | |
| H37Rv-B | SAMN32734018 | SRR23080313 | | |
| H37Rv-SC (AE1028) | SAMN32734019 | SRR23080348 | | |
| H37Rv Δ*ppsD* | SAMN32734020 | SRR23080347 | | |
| H37Rv Δ*tgs1*-1 | SAMN32734021 | SRR23080346 | | |
| H37Rv Δ*tgs1*-2 | SAMN32734022 | SRR23080345 | | |
| H37Rv Δ*tgs1*-3 | SAMN32734023 | SRR23080344 | | |
| H37Rv Δ*tgs1*-4 | SAMN32734024 | SRR23080343 | | |
| H37Rv Δ*tgs1*-5 | SAMN32734025 | SRR23080342 | SRR23080341* | |
| H37Rv Δ*tgs1*-7 | SAMN32734026 | SRR23080340 | SRR23080339* | |
| H37Rv Δ*tgs1*-8 | SAMN32734027 | SRR23080337 | | |
| H37Rv Δ*tgs1*-9 | SAMN32734028 | SRR23080336 | | |
| CDC1551-A | SAMN32734029 | SRR23080335 | | |
| CDC1551-B | SAMN32734030 | SRR23080334 | | |
| CDC1551 Δ*mas* | SAMN32734031 | SRR23080333* | | |
| Erdman | SAMN32734032 | SRR23080332 | | |
| HN878 | SAMN32734033 | SRR23080331 | | |
| KZN 4207 | SAMN32734034 | SRR23080330 | | |
| CDC1551-SC (AE5005) | SAMN32734035 | SRR23080329 | | |
| Erdman-SC (AE3003) | SAMN32734036 | SRR23080328 | | |
| HN878-SC (AE8005) | SAMN32734037 | SRR23080326 | | |
| H37Rv-B col5 | SAMN32734038 | SRR23080325 | | |
| H37Rv-B col6 | SAMN32734039 | SRR23080324 | | |
| H37Rv-B col16 | SAMN32734040 | SRR23080323 | | |
| H37Rv-B col17 | SAMN32734041 | SRR23080322 | | |
| H37Rv-B col19 | SAMN32734042 | SRR23080321 | | |
| H37Rv-B col23 | SAMN32734043 | SRR23080320 | | |
| H37Rv-B col36 | SAMN32734044 | SRR23080319 | | |

*Data acquired using Illumina MiSeq with a 300-cycle kit (76 bp reads). †Data acquired on the Illumina NextSeq system (151 bp reads). All remaining data was acquired using Illumina MiSeq with a 600-cycle kit (301 bp reads).
